# Single-cell landscape reveals NAMPT mediated macrophage polarization that regulate smooth muscle cell phenotypic switch in pulmonary arterial hypertension

**DOI:** 10.1101/2023.07.04.547668

**Authors:** Zuoshi Wen, Liujun Jiang, Fangcong Yu, Xiaodong Xu, Mengjia Chen, Jianing Xue, Pengwei Zhu, Zhangquan Ying, Zhoubin Li, Ting Chen

## Abstract

**Rationale:** Pulmonary arterial hypertension (PAH) is a progressive and lethal disease that leads to elevated pulmonary vascular resistance and right ventricular failure. The phenotypic switching of pulmonary arterial smooth muscle cells (SMCs) plays a crucial role in the pathological progression of PAH. However, the underlying mechanism of SMC phenotypic modulation remains unclear.

**Objectives:** We aim to provide a comprehensive understanding of SMC phenotypes and regulatory networks by analyzing hypertensive and non-diseased pulmonary arteries.

**Methods:** We performed single-cell RNA sequencing (scRNA-seq) on pulmonary arteries obtained from patients with PAH and healthy donors. This was followed by bioinformatics analyses, mouse models, and in vitro studies to construct a normal pulmonary artery atlas, characterize SMC phenotypes, investigate intercellular communication, and explore the molecular mechanisms underlying SMC phenotypic switching.

**Measurements and Main Results:** Our scRNA-seq analysis identified specific activation of vascular cells, including myofibrocytes, macrophage M2 polarization, endothelial-mesenchymal transition, and chondroid-like SMCs in healthy pulmonary arteries. In PAH pathology, there was an enhanced phenotypic switch of SMCs from contractile to fibroblast-like. Intercellular communication revealed increased M1 macrophage-SMC crosstalk in PAH, which was facilitated by NAMPT. Using a cellular co-culture system, we found that NAMPT-mediated M1 macrophage polarization induced fibroblast-like phenotypic switching in SMCs via the CCR2/CCR5 axis.

**Conclusions:** Our findings provide a comprehensive cell atlas of healthy human pulmonary arteries and demonstrate that NAMPT-driven M1 macrophage polarization plays a critical role in the fibroblast-like phenotypic switching of SMCs through CCR2/CCR5 cellular crosstalk in PAH.

## Introduction

Pulmonary arterial hypertension (PAH) is a progressive, life-threatening but incurable disease characterized by pulmonary arterioles vasculopathy and increased pulmonary vascular resistance, ultimately leading to right ventricular failure or death (1, 2) awaiting lung or combined heart-lung transplantation (3). The uncontrolled expansion of pulmonary arterial smooth muscle cells (SMCs) has been considered a crucial event in PAH progression (4). Despite of several advances, the underlying cellular and molecular mechanisms of PAH still remain unclear, resulting in a grim prognosis that is worse than multiple cancers (5).

SMCs, as a key to vascular remodeling, have heterogeneous phenotypes and remarkable ability to dynamically modulate from a contractile to a proliferative or synthetic state with expression of non-contractile proteins such as fibronectin and vimentin, especially in PAH (6–10). Inflammation is another major contributor to PAH, macrophages and SMCs partnered to promote migration and proliferation through key chemokine systems (11). Nevertheless, the mechanism by which macrophages interact with SMCs in pulmonary artery remains unclear.

Nicotinamide phosphoribosyltransferase (NAMPT), also known as VISFATIN or pre-B cell colony-enhancing factor (PBEF), is an enzyme involved in the synthesis of nicotinamide adenine dinucleotide (NAD+) and has been implicated in vascular remodeling process of cardiovascular diseases including atherosclerosis, hypertension and PAH (12–14). Recent studies have shown that NAMPT regulated endothelial cell and SMC activation in pulmonary vascular remodeling in PAH patients (15, 16), but it lacks further investigation to fully depict the underlying mechanisms.

In this study, by single-cell RNA sequencing (scRNA-seq) of hypertensive and non-diseased human pulmonary arteries, we firstly observed fibroblast activation, macrophage M2 polarization, endothelial-mesenchymal transition and SMC modulation in healthy pulmonary artery. Comparative analyses revealed an enhanced phenotypic switching of SMCs from contractile to fibroblast-like in neointima under PAH. Analysis of cellular communication exhibited the crucial role of pro-inflammatory macrophage-SMC crosstalk in PAH and identified NAMPT as the promoter of macrophage M1 polarization. Moreover, we generated a cellular co-culture system to reveal that NAMPT mediated macrophage M1 polarization conducted a fibroblast-like SMC phenotypic switching via CCR2/CCR5-NFκB signaling axis.

## Materials and Methods

A detailed description of the methods is provided in an online data supplement.

## Human tissue collection

The human pulmonary artery samples were collected from patients diagnosed with end-stage lung disease prior to lung transplantation, pulmonary arterial hypertension was confirmed by pre-operative echocardiogram. Healthy arteries were collected from corresponding donors during the surgery. Written informed consent was obtained from the patients. All samples were collected from the First Affiliated Hospital of Zhejiang University (China) between May 2021 and May 2022. The baseline characteristics of patients are summarized in Table E1 and E2.

## Ethics statement

The collection of human samples was approved by the research ethics committees of the Research Ethics Committees of the First Affiliated Hospital of Zhejiang University (Approval Reference No. 2021/330; Date, May 10, 2021). All procedures had local ethical approval, and all experiments were conducted according to the 2013 Declaration of Helsinki (17). All animal procedures conformed to the Guide for Care and Use of Laboratory Animals published by the US National Institutes of Health (NIH; 8th edition, 2011) and were approved by the Research Ethics Committees of the First Affiliated Hospital of Zhejiang University (Approval Reference No. 2021/159; Date, February 4, 2021). The evaluation of physiological parameters of experimental animals is shown in Table E3.

## Results

### Specific activation of vascular cells in non-diseased pulmonary artery identified by scRNA-seq

We collected hypertensive pulmonary arteries and corresponding non-diseased donor arteries during lung transplantation. Samples were subjected to scRNA-seq and histological examination to dissect cellular and molecular changes in pulmonary arterial hypertension (PAH, Figure 1A).

**Figure 1.**
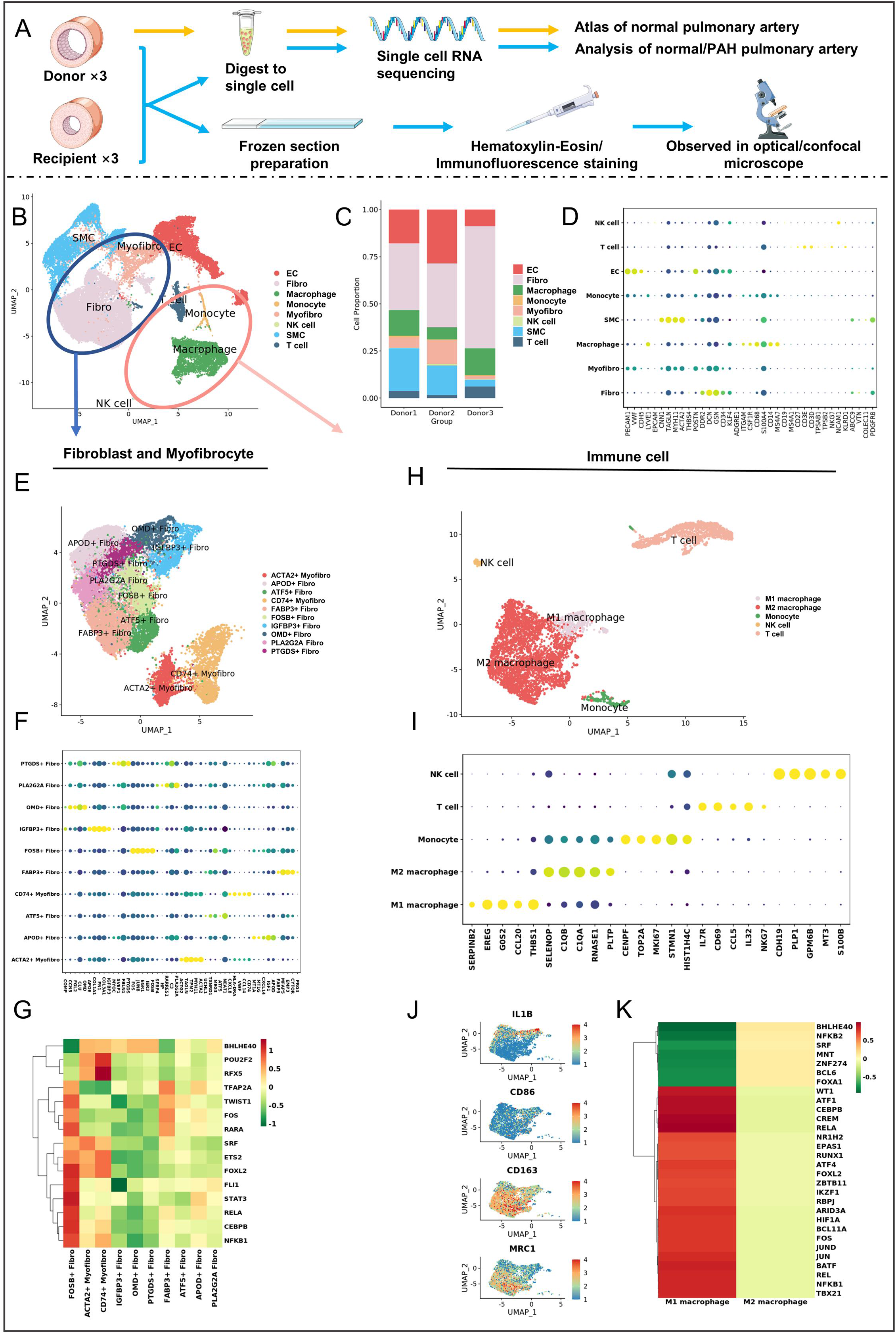
Experimental design and analysis of donor fibroblast and immune cell subpopulations. (A) Schematic representation of the experimental setup involving the collection of pulmonary artery tissue from lung transplantation donors and recipients for scRNA-seq analysis (n = 3 per group). Donor cells were used to construct a non-diseased pulmonary artery atlas, while recipient cells were compared to investigate cellular heterogeneity and dynamics in pulmonary artery hypertension. HE and immunofluorescence staining were performed to validate the observed results. **(B)** Representative UMAP plot illustrating the cell types isolated from donors. A total of 35,381 cells were pooled, and eight cell types were identified: EC (endothelial cell), Fibro (fibroblast), Macrophage, Monocyte, Myofibro (myofibrocyte), NK cell, SMC (smooth muscle cell), and T cell. Fibroblasts and myofibroblasts (highlighted by a blue circle) were subjected to dimensionality reduction clustering and UMAP visualization in **(E)**, while immune cells (enclosed by a red circle) were further displayed in **(H)**. **(C)** Bar graph displaying the cell state proportion of donors. The number of cells sequenced for each donor was as follows: Donor1, 15,295 cells; Donor2, 11,693 cells; Donor3, 8,393 cells. **(D)** Dot plot showcasing classical marker genes for cell clusters. Each dot represents a cell type, with dot size indicating the percentage of cells expressing the gene and color indicating the average expression level. **(F)** Dot plot presenting the top 5 markers for each subpopulation of fibroblasts and myofibrocytes. **(G)** Heatmap of transcription factor (TF) activities, analyzed using DoRothEA, displaying the top 15 most variable TFs across 10 individual cell clusters of fibroblasts and myofibrocytes. **(I)** Dot plot revealing the top 5 markers for each subpopulation of immune cells. **(J)** Marker genes used for cell definition, including classical markers for M1 macrophages (IL1B and CD86) and M1 macrophages (CD163 and MRC1). **(K)** Heatmap displaying the transcription fact

As a seldomly investigated tissue-type, we firstly explored the composition of normal pulmonary arteries. A total of 35,142 cells were divided into 6 major cell types (stromal: *PECAM1*^+^ endothelial cells, *MYH11*^+^ smooth muscle cells, *DCN*^+^ fibroblasts, and immune: *CD68*^+^ monocytes/macrophages, *CD3D*^+^ T cells, *NCAM1*^+^ NK cells) according to reliable type-specific gene markers. Each cell type was able to be characterized in all 3 donors despite of slight individual differences (Figure 1B-D).

Fibroblast was the dominant cell type (∼42%) in the pulmonary artery, which was further clustered into 8 fibroblast sub-clusters and 2 myofibrocyte sub-clusters according to their diverse gene expressing patterns. The cluster-defining differentially expressed genes (DEGs), transcription factors, gene ontology (GO) and Kyoto encyclopedia of genes and genomes (KEGG) pathways indicated distinct cellular function among fibroblasts, that *FABP3*^+^, *APOD*^+^ and *PLA2G2A*^+^ fibroblasts were related to lipid metabolism, OMD^+^ sub-cluster was a bone metabolism-associated one, while *FOSB*^+^ fibroblast played a role in stress response by upregulating MAPK, JAK/STAT, NF-κB signaling related genes (Figure 1E-F; Figure E1A-B; Table E4). The IGFBP3^+^ fibroblasts might be an intermediate state of myofibrocyte activation since they shared a closer relationship to myofibrocytes and were enriched in vascular smooth muscle contraction (Figure E1A-B). Myofibrocytes, which were surprisingly found even in normal arteries, were characterized by additional presence of contractile proteins and actin filaments (Figure E1C) and were divided into *ACTA2*^+^ and *CD74*^+^ sub-clusters. The *CD74*^+^ myofibrocytes markedly expressed inflammatory genes including *CXCL8*, *CCL14*, and *CD74*, that response to stress and hypoxia. The *ACTA2*^+^ myofibrocytes upregulated contraction-related genes such as *ACTG2*, *TAGLN*, and *TPM2*, which may play a role in diseases like aortic dissection/aneurysm and PAH. It was interesting that they also response to temperature stimuli and regulated epithelial cell proliferation (Figure 1E; Figure E1A-B). The transcription factor patterns between myofibrocyte sub-clusters were broadly close, except that *CD74*^+^ myofibrocytes exhibited higher enrichment of *RFX5* and *POU2F2*, that were involved in antigen presentation or regulation of T cell development (18, 19), respectively (Figure 1F).

### Macrophage polarization exists in pulmonary artery

We then further characterize 4 major immune cell types in non-diseased pulmonary artery, and found that myeloid cells especially macrophages (∼71.7%) predominate the immune populations, which could be further classified into M1 and M2 macrophages (Figure 1H-I). The M1 macrophages were characterized by *IL1B*, and expressed transcription factors including *JUN*, *FOS*, *NFKB1* and *RUNX1*, which was consistent with the pro-inflammatory potential. Conversely, the M2 macrophages, which accounted for 89.1% in macrophage population, were marked as *CD163*^+^ and *MRC1* (CD206)^+^, co-expressing transcription factors such as *ZNF274*, *FOXA1*, *BCL6* and *FOXA1* to possess anti-inflammatory and tissue repair properties (20) (Figure 1J-K; Figure E1D; Table E5).

### Unique endothelial-mesenchymal transition in pulmonary artery

Endothelial cells (ECs) represented the second largest population (∼19.2%) in normal pulmonary arteries. Although finer clustering identified 8 EC sub-clusters based on DEGs, what’s more interesting was that beyond quiescent ones there resided ECs undergoing endothelial-mesenchymal transition (EndMT ECs), even in non-diseased state (Figure 2A; Figure E2A; Table E6). EndMT ECs were characterized by upregulated contractile or fibrotic genes (*ACTA2*, *TAGLN*, *TPM2* or *DCN*, *LUM*, *COL1A2*) among canonical EC markers. Unlike quiescent ECs sharing endothelium development-related gene pattern (*CCL14*, *ACKR1*, *STC1*, *PLVAP*), EndMT ECs highly expressed genes that involved in contraction and extracellular matrix organization. Transcriptional analysis revealed overexpression of *SMAD3* and *SRF*, which indicated the crucial role of TGF-β signaling in EndMT ECs. On the other hand, downregulated functions of EndMT ECs such as antigen presentation and response to IFN-γ suggested that immune interfering might be a target for regulating the process (Figure 2B-E; Figure E2A). The phenotypic switch from quiescent to EndMT ECs was also examined by pseudotiming analysis. EndMT ECs were distributed at the terminal stage of the differentiation trajectory (Figure 2F), with downregulation of pan-EC markers and continuously increased expression of mesenchymal genes (Figure 2G; Figure E2B). Notably, chemokine-related genes (*CCL14*, *AQP1*) were found decreased in EndMT ECs, whereas genes such as *SERPINF1*, known to regulate angiogenesis (21), and *AEBP1*, involved in adipocyte metabolism (22), were upregulated across the pseudotime trajectory (Figure 2G). These results suggested a loss of inflammatory response and gain of extracellular structure organization during EndMT.

**Figure 2.**
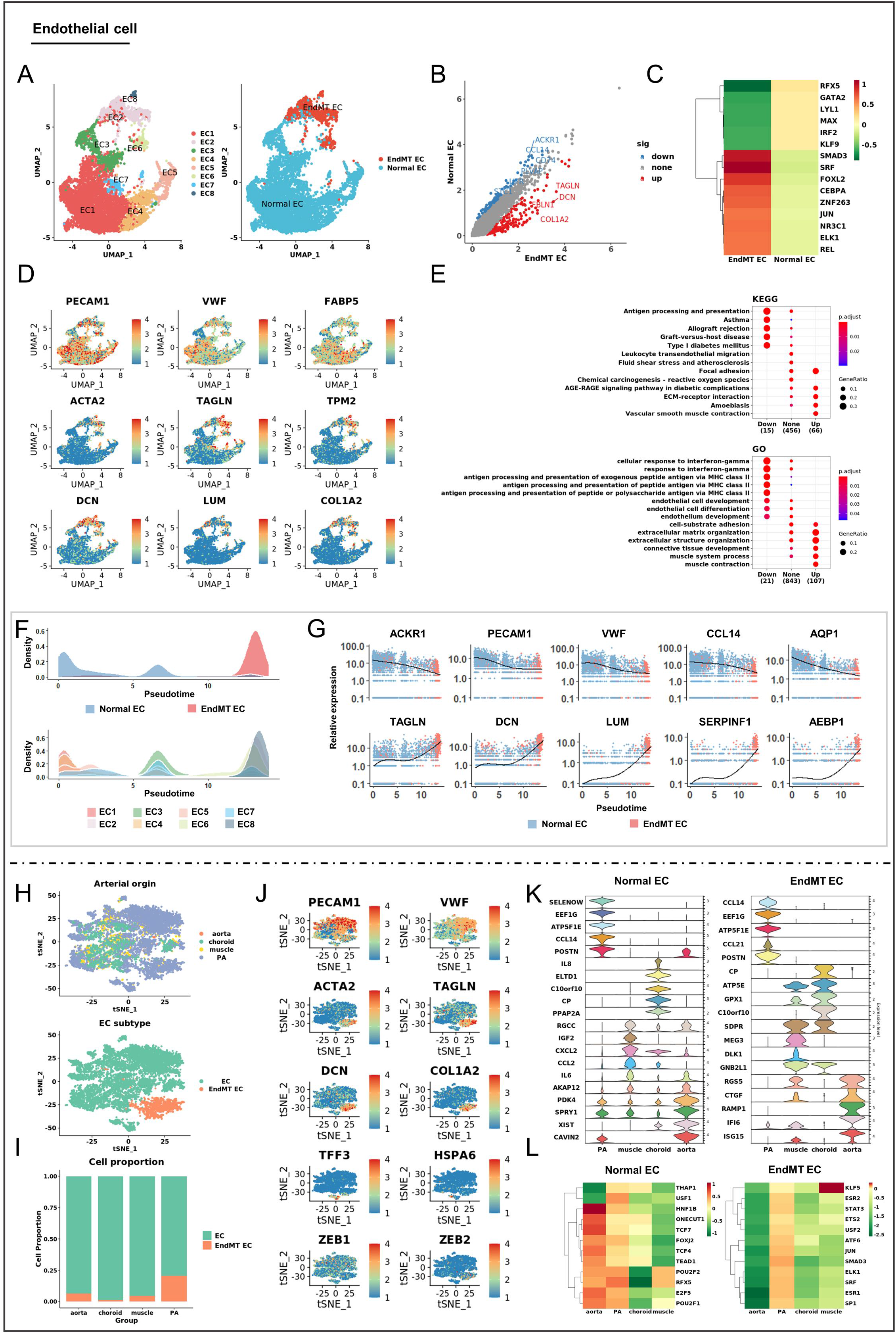
Single-cell analysis of human pulmonary artery endothelial cells. (A) UMAP plot displaying the categorization of endothelial cells into eight distinct subgroups (EC1-EC8) (left panel). Based on their transcriptomic characteristics, these subgroups were further classified as normal endothelial cells (Normal ECs) and endothelial cells undergoing endothelial-to-mesenchymal transition (EndMT ECs) (right panel). **(B)** Scatter plot illustrating the differentially expressed genes between EndMT EC and normal EC, with red dots representing upregulated genes in EndMT EC compared to Normal EC, and blue dots indicating downregulated genes. **(C)** Heatmap representing the transcription factor (TF) activities in EndMT EC and Normal EC. **(D)** Marker genes used for cell definition, including classical markers for endothelial cells (first row) and mesenchymal markers (second and third rows). **(E)** Dot plot visualization of functional enrichment analysis using KEGG and Gene Ontology (GO) terms, highlighting top differentially enriched pathways based on the upregulated and downregulated genes in EndMT EC compared to Normal EC. **(F)** Density plot displaying the distribution of endothelial cell populations, including cell types (upper panel) and cell subpopulations (lower panel), along the pseudotime trajectory. **(G)** Expression trend of selected genes along the pseudotime line. **(H)** t-SNE plot of endothelial cells derived from multiple tissues, color-coded by origin (upper panel) and cell types (lower panel). **(I)** Comparison of cell proportions in each tissues between normal and EndMT ECs. **(J)** t-SNE plots showing the expression of canonical genes for normal and EndMT ECs. **(K)** Violin plot demonstrating the expression of the top 5 differentially expressed genes for normal EC (left panel) and EndMT EC (right panel) across all tissues. **(L)** Heatmap displaying the transcription factor (TF) activities of normal EC (left panel) and EndMT EC (right panel) across all tissues.

The EndMT of ECs was then found unique in pulmonary artery. A total of 12,201 ECs from aorta (23), choroid (24), muscle (25), and pulmonary artery were collected and characterized into quiescent and EndMT ECs (Figure 2H; Figure E3). Pulmonary artery prominently consisted of 20.7% EndMT ECs, comparing to that of 6.5% in aorta, 4.5% in muscular and 1.1% in choroidal artery (Figure 2I), with more active functions of extracellular matrix organization (Figure E4). Additionally, we analyzed the expression of *ZEB* gene family, which was known a key to EndMT (26). Interestingly, *ZEB2* was more specific in arterial EndMT ECs (Figure 2J). We also identified several artery-specific genes such as *CCL14*, *CCL21*, *POSTN*, *EEF1G* for pulmonary artery, *ISG15*, *IFI6*, *RGS5*, *CTGF*, *RAMP1* for aorta, *MEG3*, *DLK1* for muscular artery and *CP*, *GPX1*, *C10orf10* for choroidal artery (Figure 2K). EndMT ECs in pulmonary artery exhibited highest transcriptional activity expressing *STAT3*, *JUN*, *SMAD3*, *ATF6*, *ELK1*, *SRF,* and *ESR1*, while *KLF5* might be vital for muscular EndMT ECs (Figure 2L).

### Modulated smooth muscle cell phenotypes in pulmonary artery

Pulmonary arterial SMCs were further split into 8 sub-clusters (Figure 3A). Using established phenotype-associated markers including contractile (*MYH11*, *CNN1*), fibrotic (*DCN*) and chondroid (*BMP2*), we categorized all cells into contractile SMCs, fibroblast-like SMCs and chondroid-like SMCs (Figure 3B-C; Figure E5; Table E7). This phenotypic switch was also examined by transcriptional or enrichment analysis showing distinct molecular and cellular functions of chondroid-like SMCs and fibroblast-like SMCs. The enrichment of *JUN*, *RELB*, *STAT1*, *NFKB1* and *SMAD3* in phenotypically switched SMCs also indicated the participation of JAK/STAT, NF-κB or PI3K/Akt signaling (Figure 3D-E).

**Figure 3.**
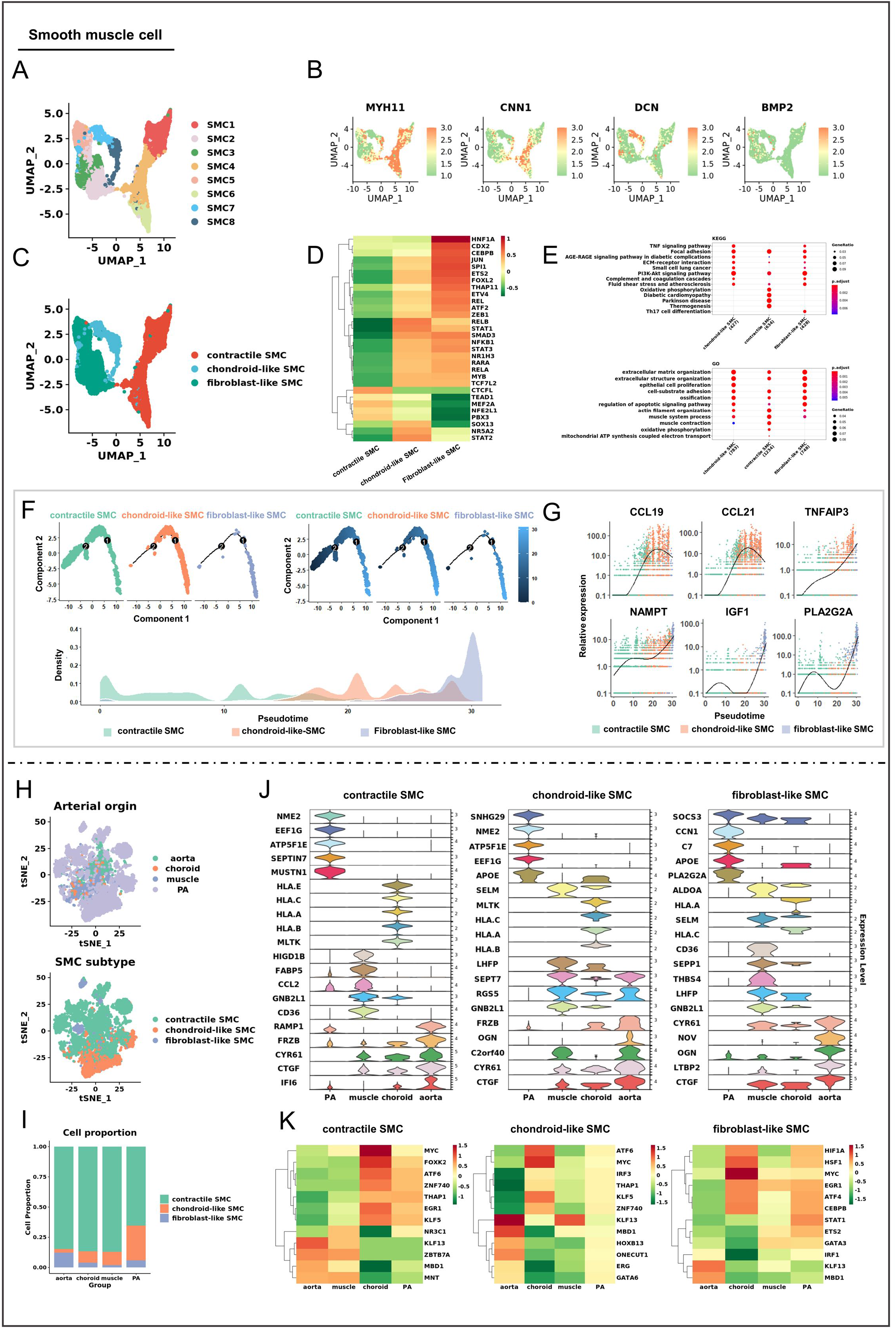
Heterogeneity of smooth muscle cells in human pulmonary artery. (A) UMAP plot categorizing smooth muscle cells into eight distinct subgroups (SMC1-SMC8). **(B)** Feature plots showing expression distribution of selected genes representing the contractile phenotype (MYH11 and CNN1), fibroblast-like phenotype (DCN), and chondroid-like phenotype (BMP2). Expression levels are color-coded and overlaid onto the UMAP plot. **(C)** Classification of smooth muscle cells based on transcriptomic characteristics into contractile smooth muscle cells (SMCs), chondroid-like smooth muscle cells (C-SMCs), and fibroblast-like smooth muscle cells (F-SMCs) in UMAP plot. **(D)** Heatmap displaying the transcription factor (TF) activities of contractile SMC, chondroid-like SMC, and fibroblast-like SMC. **(E)** Dot plot visualization of functional enrichment analysis using KEGG and Gene Ontology (GO) terms, highlighting the top differentially enriched pathways based on the differentially expressed genes of contractile SMC, chondroid-like SMC, and fibroblast-like SMC. **(F)** Pseudotime trajectory analysis of smooth muscle cell subpopulations, color-coded by cell type (upper-left panel) and pseudotime (upper-right panel). Density plot displaying the distribution of all smooth muscle cells (lower panel). **(G)** Expression trend of selected genes along the pseudotime line. **(H)** t-SNE plot of smooth muscle cells derived from multiple tissues, color-coded by origin (upper panel) and cell types (lower panel). **(I)** The bar plot showed the comparison of the proportion of all smooth muscle cells in each tissues. **(J)** Violin plot demonstrating the expression of the top 5 differentially expressed genes for contractile SMC, chondroid-like SMC, and fibroblast-like SMC across all tissues. **(K)** Heatmap displaying the transcription factor (TF) activities of contractile SMC, chondroid-like SMC, and fibroblast-like SMC across all tissues.

By tracing the transitional trajectory of SMCs, we found fibroblast-like SMCs were distributed at the terminal state, whereas chondroid-like SMCs acted as intermediate state across phenotype switching (Figure 3F). Chemokines including *CCL19* and *CCL21* conducted the transition from contractile SMCs to chondroid-like SMCs, that were downregulated during the fibroblast-like SMC transition. Intriguingly, the persistent elevation of NAMPT expression suggested an important role of fibroblast-like SMC switching (Figure 3G).

The unique pulmonary arterial SMC composition was also identified in arterial comparative analysis. The pulmonary artery possessed most chondroid-like SMCs (28.8%) while the aorta consisted of most fibroblast-like SMCs (12.2%) in non-diseased state (Figure 3H-I; Figure E6A-B). The high expression of *ATP5F1E*, *NME2*, *EEF1G* and *PLA2G2A* marked an artery-specific gene expression pattern in pulmonary arterial SMCs (Figure 3J). Transcriptional and enrichment analyses revealed *MYC*, *ATF6* and *HIF1A* involvement in unique pulmonary arterial SMCs that were related to stress response, oxidative metabolism and PAH pathology (Figure 3K; Figure E6C).

### Fibroblast-like phenotypic switch of neointimal SMCs in PAH

SMCs are capable of undergoing phenotypic switch between a contractile and a synthetic phenotype in response to various environmental cues and stimuli (10). We next sought to profile the cellular and molecular features of SMC phenotype switching under PAH. Datasets pf hypertensive and non-diseased pulmonary arteries were integrated and 12 major cell types were categorized consisting of contractile SMCs, fibroblast-like SMCs and chondroid-like SMCs (Figure 4 A). A significant increase of fibroblast-like SMCs and decrease of chondroid-like SMCs were observed in PAH (Figure 4B). Comparative analyses revealed diverse gene expression patterns that more active pro-inflammatory and pro-contractile functions in hypertensive arterial contractile SMCs and fibroblast-like SMCs via upregulation of genes including *IGFBP2*, *XIST*, *TNFRSF12A*, *JUNB*, *ACTA2*, *ACTC1* and *RGS5* (Figure 4C; E7A; Table E8). Pseudotime trajectory also revealed the phenotypic switching from contractile SMCs to fibroblast-like SMCs/chondroid-like SMCs, in which an enhanced propensity towards fibroblast-like SMCs could be observed in the course of PAH (Figure 4D). Expression levels of cytokines along the trajectory further indicated the loss of contractile phenotype, the acquisition of fibroblast-like or stressed phenotypes, and activation of NFκB and TGF-β/SMAD signaling contributed to the fibroblast-like phenotypic switch (Figure 4E; Figure E7B-C).

**Figure 4.**
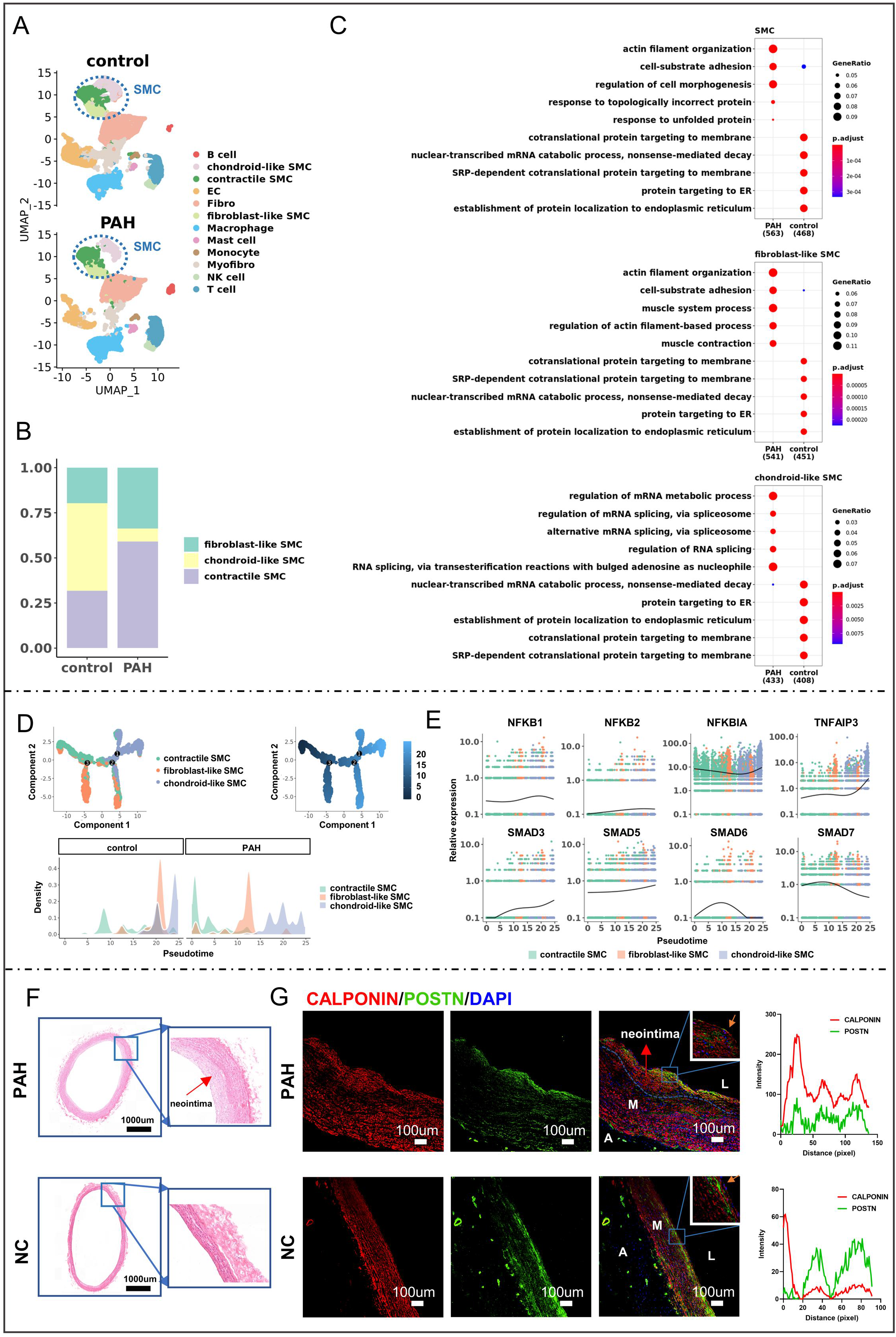
Smooth muscle cell phenotypic switching in pulmonary artery hypertension. (A) UMAP visualization of pulmonary artery cells from the normal group (upper panel) and PAH group (lower panel). **(B)** Comparison of smooth muscle cell subpopulation proportions in normal and hypertensive arteries. **(C)** Dot plot illustrating functional enrichment analysis results using Gene Ontology (GO) terms, highlighting significantly enriched pathways based on differentially expressed genes of contractile SMC, chondroid-like SMC, and fibroblast-like SMC between the control and PAH groups. **(D)** Pseudotime trajectory analysis of smooth muscle cell subpopulations, color-coded by cell type (upper-left panel) and pseudotime (upper-right panel). Density plot (lower panel) depicting the distribution of all smooth muscle cells in the control and PAH groups. **(E)** Expression profiles of selected genes along the pseudotime trajectory. **(F)** HE staining of normal and hypertensive arteries (scale bar, 1 mm), with magnification of the region highlighted in blue. The neointima in hypertensive arteries is indicated by a red arrow. **(G)** Representative immunofluorescence staining of normal and hypertensive arteries (scale bar, 100 µm). The arteries are marked with L (lumen), M (media), and A (adventitia) to identify the vascular structure, with magnification of the region highlighted in blue. The blue dashed line and red arrows roughly indicate the neointima of the pulmonary artery. Intensity trace is plotted on the right, indicating the co-localization pattern in the region marked by the orange arrow.

Further histological and immunofluorescence staining showed a remarkable neointimal hyperplasia and CALPONIN^+^ POSTN^+^ fibroblast-like SMCs within the neointima of hypertensive pulmonary arteries, particularly in the innermost layer adjacent to the lumen, suggesting a vascular remodeling involvement, which were scarcely deteced in a normal environment (Figure 4F-G). These findings indicated that in PAH, arterial SMCs exhibit a fibroblast-like phenotypic transition within the neointima, which might promote lumen obstruction.

### Macrophage M1 polarization mediated by NAMPT contributes to SMC phenotypic switch

As an active player of SMC phenotypic switch in most vascular diseases, we next investigated the changes on inflammatory patterns via examination on various ligand-receptor pairs. The cellular interactions were significantly activated in PAH, especially the inflammatory crosstalk among macrophages and contractile SMCs/fibroblast-like SMCs, which exhibited significant upregulation of incoming and outgoing interaction strengths, underscoring their pivotal role in disease progression. Conversely, chondroid-like SMCs displayed relatively inactive characteristics (Figure 5A; Figure E8A). Furthermore, macrophages of PAH exhibited a markedly elevated expression of pro-inflammatory pathway patterns, including CCL, TGF-β, TNF, IL1, IL6, and IL16, as well as growth factor regulatory pathways such as FGF, VEGF, PDGF, IGF, EGF, and HGF (Figure 5B; Figure E8C).

**Figure 5.**
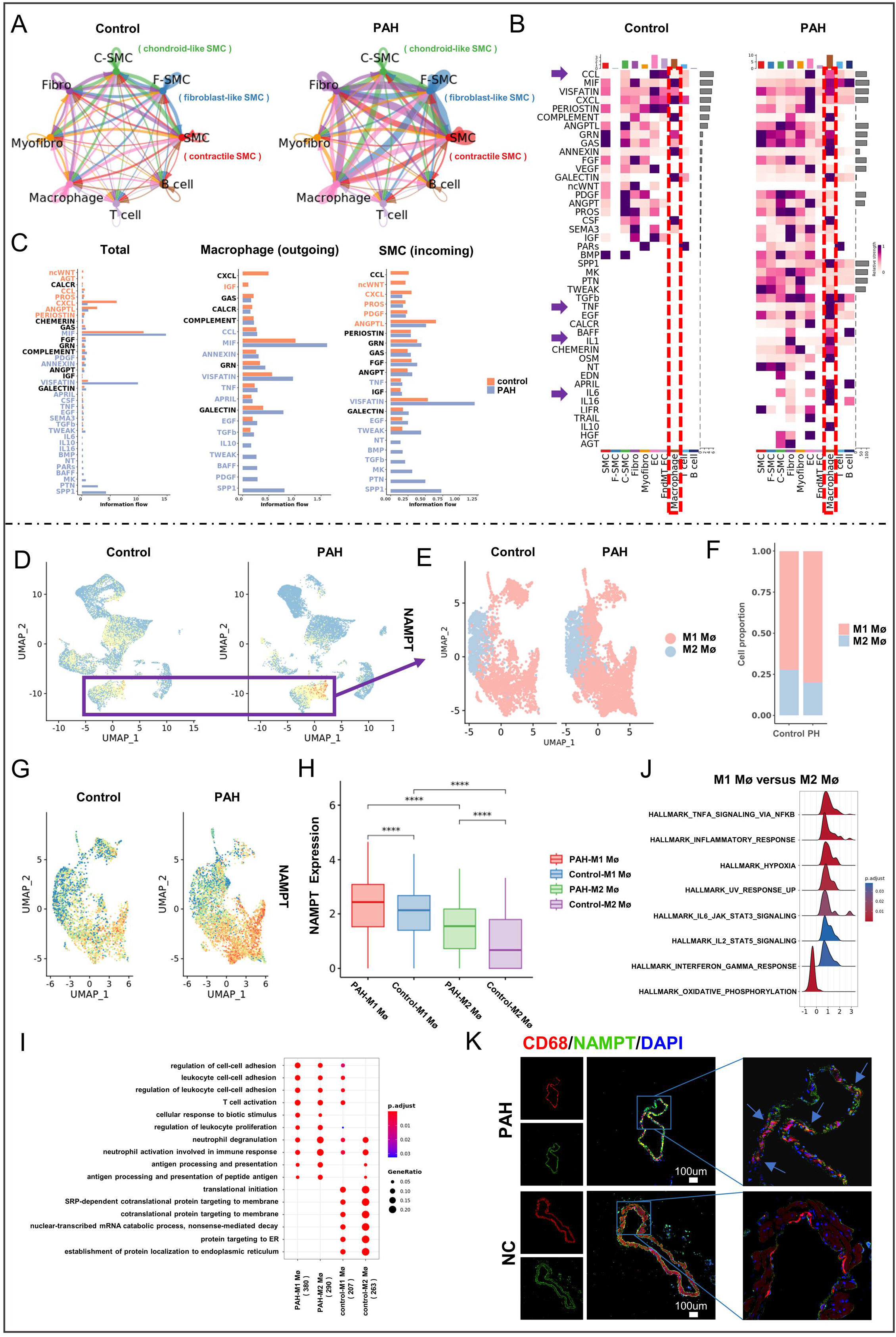
Upregulation of NAMPT expression and M1 polarization of macrophages in PAH. (A) Circle plots displaying the numbers and strengths of interactions among the main eight cell types in the normal and PAH groups. **(B)** Heatmap depicting the relative strength of signal pathway networks for each cluster, with the macrophage signaling pattern highlighted within the red dashed box. Specific signaling pathways (CCL, TNF, IL1, IL6, etc.) are indicated by purple arrows on the left. **(C)** Ranking of signaling pathways in the normal and PAH groups based on overall, macrophage output, and smooth muscle cell input information flow density. **(D)** UMAP visualization of NAMPT expression in pulmonary artery cells for the normal and PAH groups. Macrophages within the outlined box were selected for further analysis in (E). **(E)** UMAP representation of macrophage subtypes in the normal and PAH groups, categorized as M1 (red) or M2 (blue) based on marker expression. **(F)** Bar graph illustrating the proportion of M1 and M2 macrophages in normal and PAH arteries. **(G)** UMAP visualization of NAMPT expression in macrophages for the normal and PAH groups. **(H)** Boxplot displaying the NAMPT expression levels of M1 and M2 macrophages in normal and PAH groups. **(I)** Dot plot demonstrating functional enrichment analysis using Gene Ontology (GO) terms, highlighting significantly enriched pathways based on differentially expressed genes of M1 and M2 macrophages in normal and PAH groups. **(J)** Ridge plot presenting the results of GSEA functional enrichment analysis for M1 and M2 macrophages. Each peak in the plot corresponds to a gene set, with its position along the x-axis representing the enrichment score. The y-axis indicates the significance level or the running enrichment score. **(K)** Representative immunofluorescence staining of control and sugen+hypoxia murine arteries (scale bar, 100 µm). Co-stained cells are highlighted by blue arrows.

Interestingly, VISFATIN, also known as NAMPT, a pro-inflammatory marker of adipose tissue associated with systemic insulin resistance and hyperlipidemia, was predominantly enhanced in information flow intensity in the PAH group, which was especially enriched in the macrophage-SMC crosstalk (Figure 5C; Figure E8B). The upregulation of *NAMPT* was examined in the macrophage population between control and PAH groups, which was in accord with an increase of macrophage M1 polarization (Figure 5D-G) that was characterized using canonical M1/M2 markers and cellular functions (Figure 5I; Figure E8D). The elevated *NAMPT* expression in enhanced macrophage M1 polarization were further confirmed in the fractional analysis (Figure 5H), and immunofluorescence staining on murine pulmonary arteries under a Sugen (Su) plus hypoxia (Hx)-induced PAH model (Figure 5K; Figure E9A). Further enrichment analysis revealed a potential role of NFκB/TNFα signaling in M1 macrophages (Figure 5J). In summary, our results indicated that upregulation of *NAMPT* promoted the macrophages M1 polarization, that might participate in fibroblast-like SMC phenotypic switching in PAH.

### Macrophage NAMPT-SMC CCR2/CCR5 axis regulates fibroblast-like SMC phenotypic switch

To explore the impact of NAMPT on the macrophage-SMC crosstalk and the alteration of SMC biology, we established a co-culture system that composed of primary murine alveolar macrophages (upper chamber) and pulmonary arterial SMCs (lower chamber). *NAMPT* interference was performed on the upper macrophages (Figure 6A). Successful knockdown of NAMPT in macrophages following *NAMPT* small interfering RNA application (Figure 6B, 6I). We next evaluated the modulation of SMCs and observed a significant downregulation of phenotypic switching markers POSTN and RUNX2 under *NAMPT* interference (Figure 6B, 6D-E; Figure E10A), as well as a decrease migration capacity by transwell assay (Figure 6C), indicating that suppression of NAMPT in macrophages attenuated the transition of contractile SMCs to a fibroblast-like phenotype.

**Figure 6.**
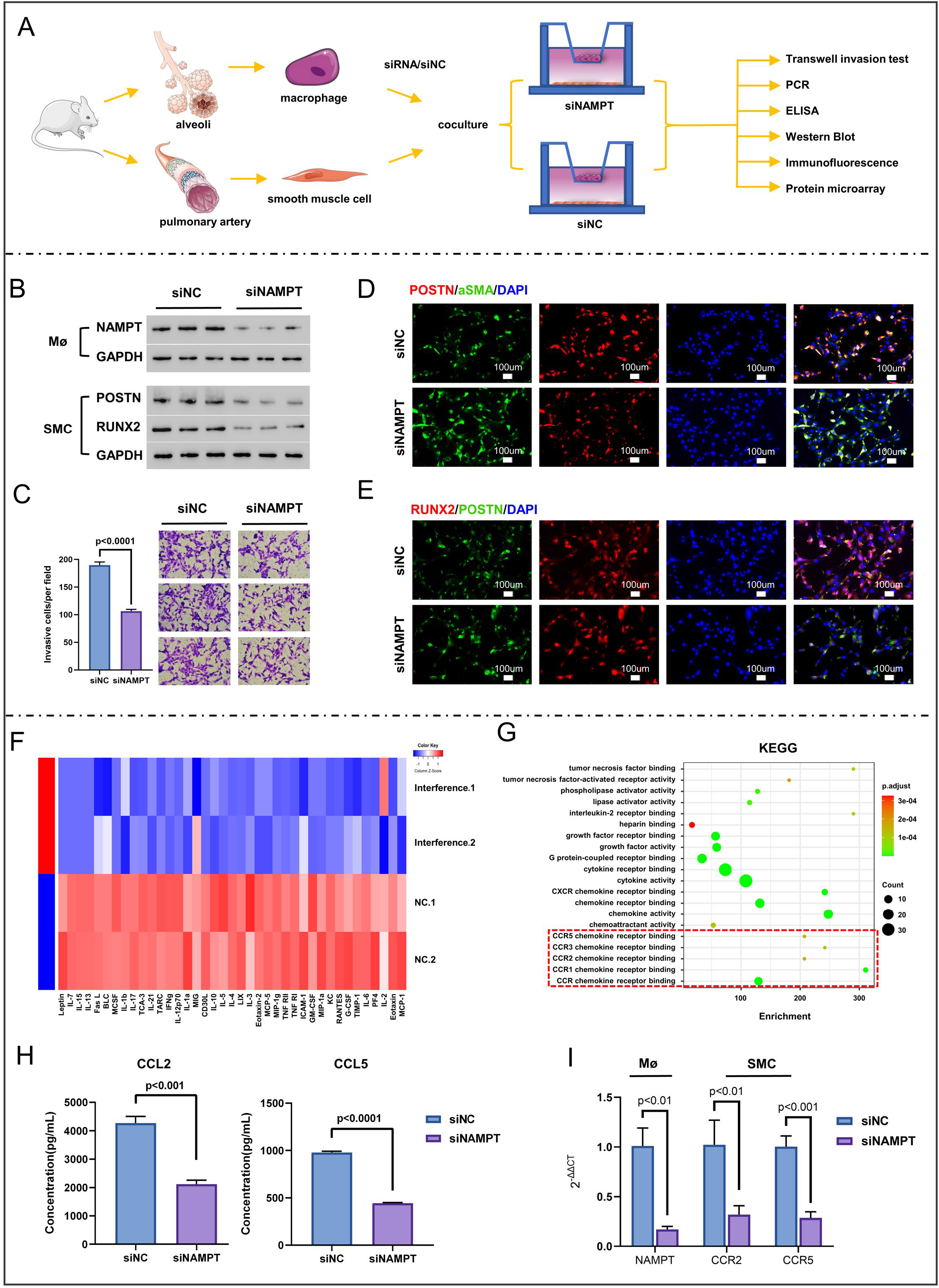
N**A**MPT**-driven macrophage polarization contributes to fibroblast-like SMC phenotypic switch (A)** A schematic representation of the experimental pipeline, demonstrating the co-culture of mouse alveolar macrophages and pulmonary artery SMCs with NAMPT interference for subsequent experiments. **(B)** Western blot analysis showing the protein expression of NAMPT in macrophage cells treated with siNC or siNAMPT, as well as the expression of phenotype switching markers (POSTN and Runx2) in SMCs. **(C)** Representative images of migrated cells stained with crystal violet (right panel), with quantification of migrated cells shown in the histogram (left panel). **(D-E)** Representative immunofluorescence staining of SMCs in the siNC and siNAMPT groups, with a scale bar of 100 µm. **(F)** A heatmap illustrating the relative enrichment of chemokines and cytokines secreted by macrophages in the co-culture system with siNAMPT or siNC, as detected by protein array analysis. Color scale: low (blue) to high (red). **(G)** A dotplot displaying the enriched pathways based on the protein array data. The size of the dots corresponds to the number of genes in each pathway, and the color of the dots represents the adjusted p-value, ranging from low (green) to high (red). The red dashed boxes indicate the pathways associated with CCR. **(H)** A bar graph showing the mean ± SD of CCL2 and CCL5 concentrations in the supernatants of the co-culture system. **(I)** A bar graph presenting the expression levels of NAMPT in macrophages and CCR2/ CCR5 in SMCs, normalized to GAPDH. The data are presented as mean ± SD from three independent experiments. Statistical significance was determined using Student’s t-test, with a p-value < 0.05 considered significant. The figure was partly generated using Servier Medical Art (http://smart.servier.com/), licensed under a Creative Commons Attribution 3.0 unported license.

To assess how NAMPT affects macrophage secretory function, cell culture supernatant from control and NAMPT knock-down macrophages were analyzed using protein microarrays. Macrophages under *NAMPT* interference downregulated the secretion of cytokines such as IL1α, IL6, IFN-γ, TNF RI and G-CSF, as well as chemokines such as MCP-1, MIP-1α and MIG (Figure 6F), which were pro-inflammatory proteins relating to cytokine activity, CCR chemokine receptor binding, response to chemokine, and cell chemotaxis (Figure 6G; Figure E9B). These results confirmed the promotion role of NAMPT in macrophage M1 polarization. Since we observed a CCR family involvement under *NAMPT* interference (Figure 6G), and macrophage might have an impact on SMC proliferation or migration via CCR2/CCR5 in PAH (11, 27, 28), we then sought to investigate the potential role of NAMPT in modulating the collaboration between macrophages and SMCs through this chemokine pattern. By ELISA analysis we quantified a significant reduction of CCL2/CCL5 under *NAMPT* interference (Figure 6H). The corresponding conspicuous downregulation of CCR2/CCR5 in SMCs was also observed (Figure 6I). The NF-κB signaling, which had been repeatedly reinforced in our study (Figure 4E, 5J), was also a critical pathway in CCL5-mediated SMC phenotype switching (29). We found that the phosphorylation of P65, an indicator of NF-κB signaling activation, was significantly attenuated under *NAMPT* interference (Figure E10B), suggesting the involvement of NF-κB signaling in NAMPT-regulated SMC phenotype switch. To conclude, under pathological PAH condition, NAMPT in macrophage promotes a M1 polarization, leads to pro-inflammatory chemokine secretion including CCL2/CCL5, that conducts a fibroblast-like SMC phenotypic switching via CCR2/CCR5-NF-κB signaling (Figure 7).

**Figure 7.**
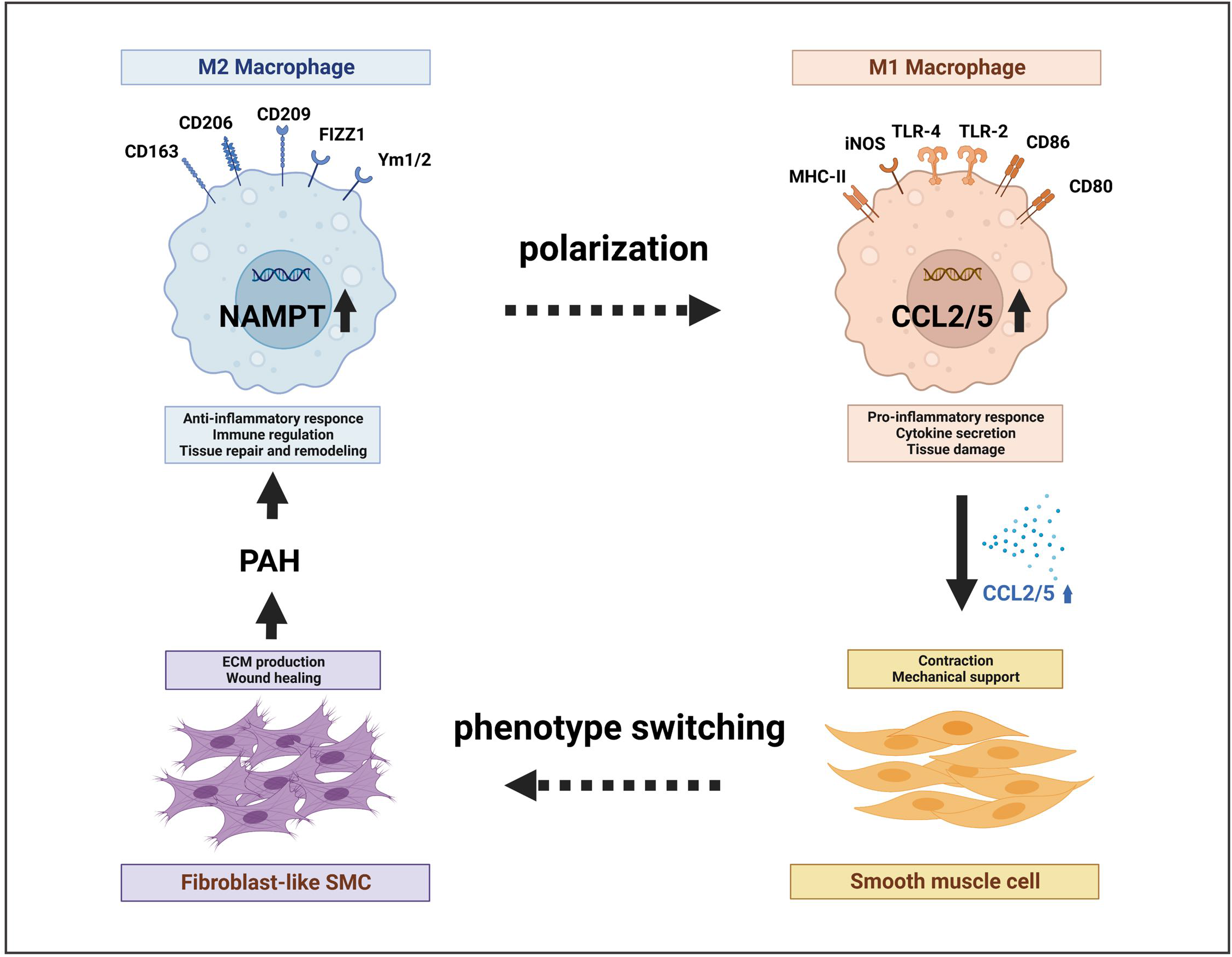
A schematic diagram illustrated the mechanism underlying the interaction of macrophage polarization and smooth muscle cell phenotypic switching in PHA.

## Discussion

Here, we utilized scRNA-seq to establish a comprehensive human cell atlas of pulmonary arteries and identified three distinct phenotypic subpopulations of pulmonary arterial SMCs: contractile, fibroblast-like, and chondroid-like. We demonstrated that the transition of SMCs from a contractile to a fibroblast-like phenotype was amplified in the neointima under PAH and was promoted by the enhanced pro-inflammatory cellular communication with macrophages. Furthermore, we identified the upregulation of NAMPT in macrophages as a key to the secretory pattern alteration. The presence of NAMPT^+^ macrophages was significantly elevated in pulmonary arteries, whereas *NAMPT* interference in macrophages effectively attenuated the M1 secretory pattern and mitigated the fibroblast-like phenotypic switching of SMCs.

The uncontrolled pulmonary vascular remodeling observed in early stages of PAH is primarily attributed to the phenotypic switching of SMCs, that characterized by excessive proliferation, migration, and invasion (10). However, the specific phenotype of SMCs that predominantly drives this process remains inadequately understood. In this study, we identified the fibroblast-like phenotype of SMCs as the primary driver of phenotypic switching in PAH, rather than chondroid-like ones. The presence of fibroblast-like SMCs within the neointima significantly contributes to lumen narrowing and elevated blood pressure. Further validation is required to elucidate the functional implications of fibroblast-like SMCs in PAH.

Macrophage polarization is a sophisticated process in response to signals from the microenvironment. Emerging evidence has suggested that macrophage balance shifts towards M1 polarization in patients with PAH (30, 31). Several studies also revealed that augmented macrophage-SMC crosstalk promotes the development of pathological SMC burden in PAH, thus garnering sustained interest among researchers (11, 32, 33). During the initial phase of inflammation, macrophages adopt an M1 phenotype and secrete pro-inflammatory molecules. However, regulating this process is essential to avoid tissue damage, and shifting towards an anti-inflammatory M2 phenotype promotes tissue healing and repair (20, 34). In atherosclerosis, apoptotic cell-clearing macrophages released factors like IL-10, TGF-β and SPMs3, which regulated the proliferation, migration and extracellular matrix production of SMCs, thereby contributed to plaque stability (35). Conversely, SMCs affected by pro-inflammatory cytokines like IL1, IL6 and TNFα released by M1 macrophages aggravated early thrombosis and stenosis of arteriovenous fistula (36). Our results revealed an increased abundance of macrophages and M1/M2 ratio in the PAH group, consistent with previous findings that an imbalanced M1/M2 ratio contributes to experimental PAH (31). Targeting the pro-inflammatory niche mediated by M1 macrophages is crucial for ameliorating vascular remodeling in PAH, as evidenced by the enrichment of signaling pathways such as IL6-JAK-STAT3, IL2-STAT5, and TNFα. Based on the theory of inflammation, experimental JAK/STAT inhibition therapy has been announced for PAH and has acquired encouraging progress (37). Other factors like MMP1 and MMP10 secreted by M1 macrophages are also involved in vascular remodeling (38).

Previous studies have revealed significant upregulation of NAMPT mRNA and protein levels in the plasma, lungs, and human lung endothelial cells of patients and rodents with PAH. This upregulation contributes to the proliferation and phenotype modulation of SMCs by enhancing store-operated calcium entry through Orai2 and STIM2 expression, as well as facilitating EndMT through paracrine signaling (15, 39–41). NAMPT exists in two forms: intracellular (iNAMPT) and extracellular (eNAMPT). iNAMPT is primarily responsible for intracellular NAD biosynthesis, while eNAMPT acts as a cytokine or adipokine to regulate systemic NAD metabolism and immune responses (12). Several studies have indicated that activation of eNAMPT secretion in pulmonary arterial endothelial cells, which was regulated by SOX and HIF-2a, promoted SMC proliferation and EndMT, ultimately leading to pulmonary vascular remodeling (15, 16). Macrophage-derived eNAMPT could sustain macrophage survival via IL-6/STAT3 signaling, thereby establishing a positive feedback loop of the effect (42). Besides macrophages, eNAMPT derived from perivascular adipose tissue exerted dose-and time-dependent effects on SMC proliferation and provided protection against oxidative stress through the activation of ERK1/2 and p38 signaling pathways (43). On the other hand, iNAMPT was notably rarely investigated in vascular diseases. It was reported to promote differentiation and polarization of tumor-associated macrophages (44). Overexpression of iNAMPT in bone marrow macrophages leads to a reduction of M1 phenotype (45). In our study, we observed increased iNAMPT expression in macrophages, aligning with findings in the diseases mentioned above. Silencing iNAMPT led to modified macrophage secretion profiles, characterized by reduced levels of pro-inflammatory molecules. This modulation attenuated the pro-inflammatory communication between macrophages and SMCs, resulting in the suppression of fibroblast-like SMC phenotype switching. In contrast to the direct effect of eNAMPT on SMC, iNAMPT exerts its influence on SMC function indirectly by regulating macrophage status and inflammatory microenvironment. Interestingly, eNAMPT has been implicated in driving M2 polarization in monocytes derived from chronic lymphocytic leukemia patients (46), which remains to be further investigated whether this forms a negative feedback regulatory mechanism with iNAMPT, contributing to the stability of macrophage status.

There are potential limitations with our work. The NAMPT mediated macrophage polarization and underlying molecular phenotypic switching process in SMCs warrant further elucidation especially in *in vivo* studies. And the effect of iNAMPT on other vascular components such as EndMT is needed for further investigation. In summary, our studies contribute valuable insights into the molecular mechanisms underlying PAH and highlight the role of NAMPT-driven macrophage polarization in promoting the fibroblast-like phenotypic switching of SMCs, suggesting iNAMPT as a promising therapeutic target for mitigating vascular remodeling and severity in PAH.

## Acknowledgements

This study was supported by the Alibaba-Zhejiang University Joint Research Center of Future Digital Healthcare and Alibaba Cloud. The authors are grateful to the Core Facility Platform of Zhejiang University School of Medicine for its technical assistance. This work was supported by a grant from the National Natural Science Foundation of China (No.82170489, 82200479); two grants from the Natural Science Foundation of Zhejiang Province (No. LR22H020001, LY23H010007); and a grant from the Project of Medical Science Research Foundation of the Health Department of Zhejiang Province (No. WKJ-ZJ-2312) and Key Laboratory of Precision Medicine for Atherosclerotic Diseases of Zhejiang Province, China(Grant No. 2022E10026)

## References

1. Ruopp NF, Cockrill BA. Diagnosis and Treatment of Pulmonary Arterial Hypertension: A Review. JAMA 2022; 327: 1379–1391.

2. Zhong L, Leng S, Alabed S, Chai P, Teo L, Ruan W, Low T-T, Wild JM, Allen JC, Lim ST, Tan JL, Yip JW-L, Swift AJ, Kiely DG, Tan R-S. Pulmonary Artery Strain Predicts Prognosis in Pulmonary Arterial Hypertension. JACC Cardiovasc Imaging 2023.

3. Hassoun PM. Pulmonary Arterial Hypertension. N Engl J Med 2021; 385: 2361–2376.

4. Thompson AAR, Lawrie A. Targeting Vascular Remodeling to Treat Pulmonary Arterial Hypertension. Trends Mol Med 2017; 23: 31–45.

5. Hoeper MM, Humbert M, Souza R, Idrees M, Kawut SM, Sliwa-Hahnle K, Jing Z-C, Gibbs JSR. A global view of pulmonary hypertension. Lancet Respir Med 2016; 4: 306–322.

6. Stenmark KR, Frid MG, Graham BB, Tuder RM. Dynamic and diverse changes in the functional properties of vascular smooth muscle cells in pulmonary hypertension. Cardiovasc Res 2018; 114: 551–564.

7. Chakraborty R, Chatterjee P, Dave JM, Ostriker AC, Greif DM, Rzucidlo EM, Martin KA. Targeting smooth muscle cell phenotypic switching in vascular disease. JVS Vasc Sci 2021; 2: 79–94.

8. Chen X, Wei X, Ma S, Xie H, Huang S, Yao M, Zhang L. Cysteine and glycine rich protein 2 exacerbates vascular fibrosis in pulmonary hypertension through the nuclear translocation of yes-associated protein and transcriptional coactivator with PDZ-binding motif. Toxicol Appl Pharmacol 2022; 457: 116319.

9. Lechartier B, Berrebeh N, Huertas A, Humbert M, Guignabert C, Tu L. Phenotypic Diversity of Vascular Smooth Muscle Cells in Pulmonary Arterial Hypertension: Implications for Therapy. Chest 2022; 161: 219–231.

10. Ma B, Cao Y, Qin J, Chen Z, Hu G, Li Q. Pulmonary artery smooth muscle cell phenotypic switching: A key event in the early stage of pulmonary artery hypertension. Drug Discov Today 2023; 28: 103559.

11. Abid S, Marcos E, Parpaleix A, Amsellem V, Breau M, Houssaini A, Vienney N, Lefevre M, Derumeaux G, Evans S, Hubeau C, Delcroix M, Quarck R, Adnot S, Lipskaia L. CCR2/CCR5-mediated macrophage-smooth muscle cell crosstalk in pulmonary hypertension. Eur Respir J 2019; 54.

12. Dahl TB, Holm S, Aukrust P, Halvorsen B. Visfatin/NAMPT: a multifaceted molecule with diverse roles in physiology and pathophysiology. Annu Rev Nutr 2012; 32: 229–243.

13. Zhou L, Zhang S, Bolor-Erdene E, Wang L, Tian D, Mei Y. NAMPT/SIRT1 Attenuate Ang II-Induced Vascular Remodeling and Vulnerability to Hypertension by Inhibiting the ROS/MAPK Pathway. Oxid Med Cell Longev 2020; 2020: 1974265.

14. Dakroub A, Nasser SA, Kobeissy F, Yassine HM, Orekhov A, Sharifi-Rad J, Iratni R, El-Yazbi AF, Eid AH. Visfatin: An emerging adipocytokine bridging the gap in the evolution of cardiovascular diseases. J Cell Physiol 2021; 236: 6282–6296.

15. Chen J, Sysol JR, Singla S, Zhao S, Yamamura A, Valdez-Jasso D, Abbasi T, Shioura KM, Sahni S, Reddy V, Sridhar A, Gao H, Torres J, Camp SM, Tang H, Ye SQ, Comhair S, Dweik R, Hassoun P, Yuan JXJ, Garcia JGN, Machado RF. Nicotinamide Phosphoribosyltransferase Promotes Pulmonary Vascular Remodeling and Is a Therapeutic Target in Pulmonary Arterial Hypertension. Circulation 2017; 135: 1532–1546.

16. Wu Y, Wharton J, Walters R, Vasilaki E, Aman J, Zhao L, Wilkins MR, Rhodes CJ. The pathophysiological role of novel pulmonary arterial hypertension gene SOX17. Eur Respir J 2021; 58.

17. Association WM. World Medical Association Declaration of Helsinki: ethical principles for medical research involving human subjects. JAMA 2013; 310: 2191–2194.

18. Corcoran L, Emslie D, Kratina T, Shi W, Hirsch S, Taubenheim N, Chevrier S. Oct2 and Obf1 as Facilitators of B:T Cell Collaboration during a Humoral Immune Response. Front Immunol 2014; 5: 108.

19. Hu Z, Zhao TV, Huang T, Ohtsuki S, Jin K, Goronzy IN, Wu B, Abdel MP, Bettencourt JW, Berry GJ, Goronzy JJ, Weyand CM. The transcription factor RFX5 coordinates antigen-presenting function and resistance to nutrient stress in synovial macrophages. Nat Metab 2022; 4: 759–774.

20. Shapouri-Moghaddam A, Mohammadian S, Vazini H, Taghadosi M, Esmaeili S-A, Mardani F, Seifi B, Mohammadi A, Afshari JT, Sahebkar A. Macrophage plasticity, polarization, and function in health and disease. J Cell Physiol 2018; 233: 6425–6440.

21. Berghoff AS, Ilhan-Mutlu A, Dinhof C, Magerle M, Hackl M, Widhalm G, Hainfellner JA, Dieckmann K, Pichler J, Hutterer M, Melchardt T, Bartsch R, Zielinski CC, Birner P, Preusser M. Differential role of angiogenesis and tumour cell proliferation in brain metastases according to primary tumour type: analysis of 639 cases. Neuropathol Appl Neurobiol 2015; 41: e41–e55.

22. Rao M, Wang X, Guo G, Wang L, Chen S, Yin P, Chen K, Chen L, Zhang Z, Chen X, Hu X, Hu S, Song J. Resolving the intertwining of inflammation and fibrosis in human heart failure at single-cell level. Basic Res Cardiol 2021; 116: 55.

23. Li Y, Ren P, Dawson A, Vasquez HG, Ageedi W, Zhang C, Luo W, Chen R, Li Y, Kim S, Lu HS, Cassis LA, Coselli JS, Daugherty A, Shen YH, LeMaire SA. Single-Cell Transcriptome Analysis Reveals Dynamic Cell Populations and Differential Gene Expression Patterns in Control and Aneurysmal Human Aortic Tissue. Circulation 2020; 142: 1374–1388.

24. Voigt AP, Mulfaul K, Mullin NK, Flamme-Wiese MJ, Giacalone JC, Stone EM, Tucker BA, Scheetz TE, Mullins RF. Single-cell transcriptomics of the human retinal pigment epithelium and choroid in health and macular degeneration. Proc Natl Acad Sci U S A 2019; 116: 24100–24107.

25. De Micheli AJ, Spector JA, Elemento O, Cosgrove BD. A reference single-cell transcriptomic atlas of human skeletal muscle tissue reveals bifurcated muscle stem cell populations. Skelet Muscle 2020; 10: 19.

26. Fu R, Li Y, Jiang N, Ren B-X, Zang C-Z, Liu L-J, Lv W-C, Li H-M, Weiss S, Li Z-Y, Lu T, Wu Z-Q. Inactivation of endothelial ZEB1 impedes tumor progression and sensitizes tumors to conventional therapies. J Clin Invest 2020; 130: 1252–1270.

27. Amsellem V, Lipskaia L, Abid S, Poupel L, Houssaini A, Quarck R, Marcos E, Mouraret N, Parpaleix A, Bobe R, Gary-Bobo G, Saker M, Dubois-Randé J-L, Gladwin MT, Norris KA, Delcroix M, Combadière C, Adnot S. CCR5 as a treatment target in pulmonary arterial hypertension. Circulation 2014; 130: 880–891.

28. Amsellem V, Abid S, Poupel L, Parpaleix A, Rodero M, Gary-Bobo G, Latiri M, Dubois-Rande J-L, Lipskaia L, Combadiere C, Adnot S. Roles for the CX3CL1/CX3CR1 and CCL2/CCR2 Chemokine Systems in Hypoxic Pulmonary Hypertension. Am J Respir Cell Mol Biol 2017; 56: 597–608.

29. Lin C-S, Hsieh P-S, Hwang L-L, Lee Y-H, Tsai S-H, Tu Y-C, Hung Y-W, Liu C-C, Chuang Y-P, Liao M-T, Chien S, Tsai M-C. The CCL5/CCR5 Axis Promotes Vascular Smooth Muscle Cell Proliferation and Atherogenic Phenotype Switching. Cell Physiol Biochem 2018; 47: 707–720.

30. Hashimoto-Kataoka T, Hosen N, Sonobe T, Arita Y, Yasui T, Masaki T, Minami M, Inagaki T, Miyagawa S, Sawa Y, Murakami M, Kumanogoh A, Yamauchi-Takihara K, Okumura M, Kishimoto T, Komuro I, Shirai M, Sakata Y, Nakaoka Y. Interleukin-6/interleukin-21 signaling axis is critical in the pathogenesis of pulmonary arterial hypertension. Proc Natl Acad Sci U S A 2015; 112: E2677–E2686.

31. Zawia A, Arnold ND, West L, Pickworth JA, Turton H, Iremonger J, Braithwaite AT, Cañedo J, Johnston SA, Thompson AAR, Miller G, Lawrie A. Altered Macrophage Polarization Induces Experimental Pulmonary Hypertension and Is Observed in Patients With Pulmonary Arterial Hypertension. Arterioscler Thromb Vasc Biol 2021; 41: 430–445.

32. Koga J-i, Aikawa M. Crosstalk between macrophages and smooth muscle cells in atherosclerotic vascular diseases. Vascul Pharmacol 2012; 57: 24–28.

33. Ntokou A, Dave JM, Kauffman AC, Sauler M, Ryu C, Hwa J, Herzog EL, Singh I, Saltzman WM, Greif DM. Macrophage-derived PDGF-B induces muscularization in murine and human pulmonary hypertension. JCI Insight 2021; 6.

34. Wculek SK, Dunphy G, Heras-Murillo I, Mastrangelo A, Sancho D. Metabolism of tissue macrophages in homeostasis and pathology. Cell Mol Immunol 2022; 19: 384–408.

35. Yurdagul A. Crosstalk Between Macrophages and Vascular Smooth Muscle Cells in Atherosclerotic Plaque Stability. Arterioscler Thromb Vasc Biol 2022; 42: 372–380.

36. Rai V, Singh H, Agrawal DK. Targeting the Crosstalk of Immune Response and Vascular Smooth Muscle Cells Phenotype Switch for Arteriovenous Fistula Maturation. Int J Mol Sci 2022; 23.

37. Yerabolu D, Weiss A, Kojonazarov B, Boehm M, Schlueter BC, Ruppert C, Günther A, Jonigk D, Grimminger F, Ghofrani H-A, Seeger W, Weissmann N, Schermuly RT. Targeting Jak-Stat Signaling in Experimental Pulmonary Hypertension. Am J Respir Cell Mol Biol 2021; 64: 100–114.

38. Chi P-L, Cheng C-C, Hung C-C, Wang M-T, Liu H-Y, Ke M-W, Shen M-C, Lin K-C, Kuo S-H, Hsieh P-P, Wann S-R, Huang W-C. MMP-10 from M1 macrophages promotes pulmonary vascular remodeling and pulmonary arterial hypertension. Int J Biol Sci 2022; 18: 331–348.

39. van der Veer E, Ho C, O’Neil C, Barbosa N, Scott R, Cregan SP, Pickering JG. Extension of human cell lifespan by nicotinamide phosphoribosyltransferase. J Biol Chem 2007; 282: 10841–10845.

40. Borradaile NM, Pickering JG. Nicotinamide phosphoribosyltransferase imparts human endothelial cells with extended replicative lifespan and enhanced angiogenic capacity in a high glucose environment. Aging Cell 2009; 8: 100–112.

41. Sun X, Sun BL, Babicheva A, Vanderpool R, Oita RC, Casanova N, Tang H, Gupta A, Lynn H, Gupta G, Rischard F, Sammani S, Kempf CL, Moreno-Vinasco L, Ahmed M, Camp SM, Wang J, Desai AA, Yuan JXJ, Garcia JGN. Direct Extracellular NAMPT Involvement in Pulmonary Hypertension and Vascular Remodeling. Transcriptional Regulation by SOX and HIF-2α. Am J Respir Cell Mol Biol 2020; 63.

42. Li Y, Zhang Y, Dorweiler B, Cui D, Wang T, Woo CW, Brunkan CS, Wolberger C, Imai S-i, Tabas I. Extracellular Nampt promotes macrophage survival via a nonenzymatic interleukin-6/STAT3 signaling mechanism. J Biol Chem 2008; 283: 34833–34843.

43. Wang P, Li W-L, Liu J-M, Miao C-Y. NAMPT and NAMPT-controlled NAD Metabolism in Vascular Repair. J Cardiovasc Pharmacol 2016; 67: 474–481.

44. Sica A, Strauss L, Consonni FM, Travelli C, Genazzani A, Porta C. Metabolic regulation of suppressive myeloid cells in cancer. Cytokine Growth Factor Rev 2017; 35: 27–35.

45. Bermudez B, Dahl TB, Medina I, Groeneweg M, Holm S, Montserrat-de la Paz S, Rousch M, Otten J, Herias V, Varela LM, Ranheim T, Yndestad A, Ortega-Gomez A, Abia R, Nagy L, Aukrust P, Muriana FJG, Halvorsen B, Biessen EAL. Leukocyte Overexpression of Intracellular NAMPT Attenuates Atherosclerosis by Regulating PPARγ-Dependent Monocyte Differentiation and Function. Arterioscler Thromb Vasc Biol 2017; 37: 1157–1167.

46. Audrito V, Serra S, Brusa D, Mazzola F, Arruga F, Vaisitti T, Coscia M, Maffei R, Rossi D, Wang T, Inghirami G, Rizzi M, Gaidano G, Garcia JGN, Wolberger C, Raffaelli N, Deaglio S. Extracellular nicotinamide phosphoribosyltransferase (NAMPT) promotes M2 macrophage polarization in chronic lymphocytic leukemia. Blood 2015; 125: 111–123.

