## Supplementary material+ Table E1,2 for "Single-cell landscape reveals NAMPT mediated macrophage polarization that regulate smooth muscle cell phenotypic switch in pulmonary arterial hypertension"

#### Online Data Supplement

**Supplementary Methods**

#### 1. Single-cell Isolation and Library Preparation

Single-cell RNA sequencing (scRNA-seq) was performed on human pulmonary arteries using established protocols[1]. Briefly, pulmonary arteries were first washed with PBS, minced into small pieces on ice, and enzymatically digested using a combination of collagenase I (Gibco, 17018029) and dispase II (Sigma, D4693) in Hank's balanced salt solution containing calcium and magnesium (HBSS, Gibco, 14025092). After digestion, the samples were sieved through a 40-µm cell strainer and centrifuged at 300g for 5 min. The red blood cell lysis buffer (Miltenyi Biotec, 130-094-183) was added re remove erythrocytes. Live cells were then sorted using a MACS Dead Cell Removal Kit (Miltenyi Biotec, 130-090-101) and re-suspended in PBS with 0.04% BSA. scRNA-seq was conducted using Chromium™ Single Cell 3' Reagent Kit v3.1 chemistry (10x Genomics) following standard procedures. Libraries were sequenced on a Novaseq6000 PE150 platform (Illumina) using a paired-end 150 bp sequencing strategy. The 10x Chromium procedure, library generation, and sequencing were performed by Oebiotech Co., Ltd (Shanghai, China).

#### 2. Single-cell RNA Sequencing Data Preprocessing

The raw scRNA-seq data was processed using 10x Genomics Cell Ranger software (version 6.1.0). The pre-processed matrix was analyzed using the Seurat suite (version 4.3.0) in R Studio (version 4.2.3)[2]. Quality control steps were performed, including the removal of doublet cells using the scDblFinder R package[3] and filtering out genes detected in less than three cells. Cells expressing fewer than 400 and more than 6000 genes, as well as those with more than 20% mitochondrial UMIs, were excluded from analysis, as they likely represent cell multiplets and stressed cells, respectively. After quality control, a total of 62,375 single cells and 27,583 genes were retained for downstream analyses. Library size normalization was conducted using the Seurat function NormalizeData, which normalized the UMI count for each cell by total expression and produced a log-transformed result multiplied by 10,000. To focus on strong biological signals, the Seurat function FindVariableFeatures was used to select the top 2000 genes with the highest cell-to-cell variation, known as variable genes. Scaling was performed using the Seurat function ScaleData to scale and center normalized counts. Batch effect correction was applied to the selected 2000 variable genes across six samples using the standard Seurat V3 integration algorithm. Anchor cells were identified from the six different samples based on mutual nearest neighbors, and the expression matrix was corrected accordingly. An integrated and batch-corrected expression matrix for all cells was obtained and used for subsequent analyses.

#### 3. Dimensional Reduction and Clustering

Principal component analysis (PCA) was conducted using the Seurat function RunPCA to reduce the dimensionality of the dataset. To determine the appropriate number of principal components (PCs), the significance of the PCs was estimated using the Seurat function JackStraw. We identified the first 30 PCs as significant and utilized them for subsequent analyses. Uniform manifold approximation and projection (UMAP) were performed using the Seurat function RunUMAP to visualize the cells in two dimensions. Prior to clustering, we constructed a shared nearest neighbor graph based on the Euclidean distance in the PCA space using the Seurat function FindNeighbors. Clustering was then carried out using the Seurat function FindClusters with the Louvain algorithm, employing a resolution of 0.5. Marker genes for each cluster were determined using the Wilcoxon rank-sum test via the Seurat FindAllMarkers function. Cluster identities were annotated by manually matching cell type-specific marker genes to the marker genes for the clusters, with the cell type-specific marker genes obtained from canonical marker genes.

#### 4. Gene Enrichment Analyses

Gene enrichment analyses were conducted using the clusterProfiler R package[4,5]. Enriched genes that were upregulated in each cluster were identified using the "FindMarkers" function (min.pct=0.25, logfc.threshold=0.25, test.use="wilcox"). The top 200 genes, ordered by fold change, were selected for subsequent gene enrichment analysis. The biological process (BP) of Gene Ontology was annotated using the "EnrichGO" function with the annotation database org.Hs.eg.db. The BP expression level refers to the average expression of the gene sets in the BP annotation, obtained from the online database (geneontology.org).

#### 5. Pseudotime Trajectory Analyses

Pseudotime trajectory analysis was conducted using the monocle R package[6–8] (version 2.14) with default settings, unless stated otherwise. Genes used for pseudotime ordering were selected from the differentially expressed genes with a q-value < 0.05, identified using the "differential gene test" function with the fullModelFormulaStri set as pseudotime. Dimension reduction and cell ordering along the pseudotime trajectory were performed using the DDRTree method. Gene expression levels at different pseudotime points were compared using the "differentialGeneTest" function, and the expression profiles of selected genes were visualized using the "plot_genes_in_pseudotime" function.

#### 6. Infer transcription factor activities

We employed the DoRothEA package[6,9,10] to evaluate the relative activity levels of transcription factors (TFs) across different cell types. DoRothEA calculates TF activity levels based on the expression levels of their target genes rather than the TFs themselves. In our analysis, we focused on highly confident interactions between TFs and target genes, specifically those with confidence levels "A," "B," or "C." We utilized VIPER for statistical analysis of TF activity levels, as VIPER considers the mode of each TF-target interaction and has been demonstrated to be suitable for single-cell analysis[10,11]. We compared TF activity levels between cell types and visualized the top TFs based on variance in activity levels.

#### 7. Cell-Cell Communication Analyses

We utilized the CellChat package[12] to analyze cell-cell communication networks in our single-cell RNA sequencing data. The communication strength between cell clusters, indicating intercellular communication probability, was assessed using the "computeCommunProb" function. Aggregated communication networks between clusters were generated by summarizing the communication probabilities. These networks were visualized in various formats using the "netVisual_circle/heatmap/bubble" function. Furthermore, we compared the information flow in each signaling pathway between the control and PAH groups using the "compareInteractions" function. By leveraging CellChat, we gained valuable insights into the complex communication patterns within our single-cell dataset.

**8. Experimental Animals**

All animals were fed a standard laboratory diet with free access to food and water and kept in a temperature- (22 °C ± 1°C) and humidity-controlled (65%-70%) room, with a 12-h light–dark cycle. All mice generated or purchased were housed in the First Affiliated Hospital of Zhejiang University School of Medicine (Zhejiang, China) for at least 1 week before use. Male C57BL/6 mice weighing 20-25 g were assigned to two groups: the Sugen5416 plus hypoxia (Su/Hx) group and the control group. In the Su/Hx group, mice received single subcutaneous injection of SU5416 (20 mg/kg/week), as well as placed in a hypoxic chamber (10% oxygen) for 3 weeks. The control group received a saline injection and was housed under normoxic conditions. Hemodynamic measurements were conducted to assess the development of pulmonary hypertension. The Sugen5416 was dissolved in CMC buffer (0.5% sodium carboxymethyl cellulose, 0.9% sodium chloride, 0.4% polysorbate-80 and 0.9% benzyl alcohol) to prepare a suspension, which was freshly prepared before use. Recombinant mouse IL-11 protein was dissolved in sterile water for injection, aliquoted as needed, and stored at -80°C.

After 3 weeks of Sugen5416 plus hypoxia exposure, mice were subjected to anesthesia using $CO_2$ according to the designated time points following the ischemic models, in accordance with the guidelines set by the NIH, while ensuring minimal stress to the animals. In brief, a cage containing three to five mice was placed within a separate chamber with a volume of 20 liters. A cylinder containing compressed 99.99% CO2 gas was connected to the chamber, and the gas was introduced at a flow rate of 10 liters per minute. Within 3 minutes, all mice became unconscious and ceased spontaneous breathing. After an additional minute of $CO_2$ flow, the mice were reevaluated to confirm the absence of respiration, as indicated by the fading of eye color and lack of pupillary response to light. The mice were then removed from the cage, placed on a heating pad in supine position, and fixed their limbs. The chest hair was shaved, and a GE Vivid E95 color Doppler ultrasound diagnostic device with a corresponding animal probe was used to perform cardiac ultrasound examination on the experimental animals, and measure the pulmonary artery VTI (Velocity Time Integral), PAT (Pulmonary artery Acceleration Time), PET (Pulmonary artery Ejection Time) and TAPSE (tricuspid annular plane systolic excursion), as well as the left ventricular ejection fraction (LVEF). The limbs and incisors were fixed, and the abdominal skin was cut open with scissors. The abdominal cavity was opened, and the diaphragm was exposed. A catheter was inserted through the diaphragm to puncture the right ventricle, and the RVSP of the experimental animals was measured according to the pressure waveform curve and pressure value changes displayed on the computer terminal. The data were recorded when the wave value was stable. The software used for data recording and analysis were PHILIPS SureSigns VM6. The pulmonary artery was collected simultaneously for subsequent experiments.

#### 9. Immunofluorescence staining

For immunofluorescence staining of frozen tissue sections (mouse/human), tissues were collected, washed with PBS, and fixed in 4% paraformaldehyde at 4 °C overnight. After fixation, tissues were dehydrated in 35% sucrose solution at 4 °C overnight. Subsequently, tissues were embedded in OCT, frozen at -80 °C for storage, or sectioned into 7-µm sections using a cryostat (Leica CM1950). The sections were then blocked and permeabilized in 5% BSA solution (containing 0.1% Triton X-100) for 1 hour. Primary antibodies were applied overnight, followed by incubation with Alexa Fluor-conjugated secondary antibodies (Invitrogen, 1:500) for 1 hour. DAPI staining was performed for nuclear visualization. Primary antibodies used in this study included CALPONIN (Abcam, ab46794, 1:200), POSTN (R&D, af2955, 1:50), CD68 (Abcam, AB125212, 1:100), and NAMPT (Santa Cruz, sc-166866, 1:100). Images were acquired using an Olympus FV3000 confocal laser scanning microscope and further analyzed with NIH-Fiji software.

#### 10. Hematoxylin-Eosin staining

Frozen tissue sections were allowed to air-dry and were subsequently fixed in 4% paraformaldehyde for 10 minutes. Following PBS washing, the sections were stained with hematoxylin solution for 5 minutes, and excess stain was removed by rinsing in tap water. The sections were then differentiated in 1% acid alcohol solution for 1 minute, followed by another rinse in tap water. To provide contrast, the sections were counterstained with eosin solution for 1 minute. After dehydration in graded ethanol solutions, the sections were cleared in xylene and mounted with a coverslip using mounting medium. The stained sections were then observed under a light microscope, and images were captured.

**11. siRNA Transfection for NAMPT Knockdown**

Mouse primary alveolar macrophages were seeded at a density of approximately 500,000 cells per well in 6-well plates one day prior to transfection. The cells reached 30-50% confluency on the following day. For transfection, the medium in each well was replaced with 2 ml of fresh serum-containing culture medium without antibiotics. Two sterile centrifuge tubes were prepared: one containing 125 μl of serum-free DMEM medium or Opti-MEM® Medium, and the other containing 100 pmol of siRNA and 5 μl of Lipo6000™ transfection reagent (Biouniquer). The contents of each tube were gently mixed by pipetting. After a 5-minute incubation at room temperature, the siRNA-containing medium was added to the tube with the Lipo6000™ transfection reagent, mixed gently, and incubated for another 5 minutes at room temperature. Next, 250 μl of the Lipo6000™ transfection reagent-siRNA mixture was evenly added to each well of the 6-well plate. The plate was gently swirled to ensure uniform distribution. The cells were incubated with the transfection mixture for 6 hours, followed by replacement with fresh complete culture medium. The transfected cells were further incubated for 48 hours under standard culture conditions.

**12. Cell Co-culture System**

Macrophages and mouse pulmonary arterial smooth muscle cells were co-cultured in a transwell system using DMEM medium supplemented with 10% fetal bovine serum and 1% penicillin-streptomycin. The co-culture was divided into two groups: Group A, which consisted of macrophages in the upper chamber (treated with IgG control), and mouse pulmonary arterial smooth muscle cells in the lower chamber; and Group B, which consisted of macrophages transfected with NAMPT interference plasmid in the upper chamber, and mouse pulmonary artery smooth muscle cells in the lower chamber.

**13. Cell Immunofluorescence staining**

Cells were fixed in 4% paraformaldehyde for 10 minutes at room temperature, followed by three washes with PBS. Subsequently, cells were incubated with 0.5% Triton X-100 (Invitrogen) for 15 minutes, blocked with PBS containing 5% bovine serum albumin for 1 hour, and then incubated with primary antibodies overnight at 4 °C. On the following day, cells were washed three times with PBS and incubated with Alexa Fluor-conjugated secondary antibodies (Invitrogen, 1:500) for 2 hours at room temperature. Nuclei were visualized by DAPI staining. Primary antibodies used in this study included POSTN (R&D, af2955, 1:50), αSMA (Sigma, F3777, 1:500), and RUNX2 (Abcam, ab76956, 1:100). The stained cells were observed and analyzed using an Olympus FV3000 confocal laser scanning microscope.

**14. Quantitative Polymerase Chain Reaction (qPCR)**

Total RNA was extracted from the samples using the TRleasyTM Total RNA Extraction Reagent (Yeasen, 10606ES60). The concentration and purity of the RNA were assessed using a spectrophotometer. Reverse transcription was performed using the Hifair® II 1st Strand cDNA Synthesis Kit (Yeasen, 11119ES60), with 1 μg of total RNA used for cDNA synthesis. The qPCR reactions were prepared using the Hieff® qPCR SYBR Green Master Mix (Yeasen, 11203ES03) and specific primers.

| | |
|---|---|
| NAMPT | Forward, 5'-TGGGGTGAAGACCTGAGACA-3' |
| | Reverse, 5'-CCACCAGAACCGAAGGAGAC-3' |
| CCR2 | Forward, 5’-ACACCCTGTTTCGCTGTAGG-3’ |
| | Reverse, 5’-TGCATGGCCTGGTCTAAGTG-3’ |

| | |
|---|---|
| CCR5 | Forward, 5'-AAGTGTAGTCACTTGGGCGG-3' |
| | Reverse, 5'-CCTACAGCGAAACAGGGTGT-3' |
| GADPH | Forward, 5'-CTGCCCAGAACATCATCC-3' |
| | Reverse, 5'-CTCAGATGCCTGCTTCAC-3' |

The primers used were as follows:

The thermal cycling conditions included an initial denaturation step at 95°C for 5 minutes, followed by 40 cycles of denaturation at 95°C for 15 seconds, annealing at 60°C for 30 seconds, and extension at 72°C for 30 seconds. Fluorescence signals were measured during the extension step of each cycle. A melting curve analysis was performed to confirm the specificity of the amplification. The relative gene expression levels were calculated using the $2^{-\Delta\Delta Ct}$ method, normalized to an internal reference gene. Each sample was run in triplicate, and the average Ct values were used for analysis.

**15. Enzyme-linked immunosorbent assay**

The levels of CCL2 and CCL5 in the culture medium were quantified using the Mouse ELISA Kit (MultiSciences EK287/2 48，EK287/2-96). Samples and standards were prepared according to the manufacturer's instructions. Scanning and data extraction were conducted by RayBiotech Inc. Cytokine concentrations in the samples were determined by comparing the signals to the standard curve, and the results were plotted using Prism Software (GraphPad).

**16. Western blot analysis**

To extract the total cellular proteins, cells were washed 2-3 times with PBS, and the residual liquid was aspirated using a pipette. RIPA lysis buffer (Thermo Scientific™, 89900) supplemented with protease inhibitors was added to the culture dish or flask and incubated with intermittent shaking for 3-5 minutes. The cells were then scraped using a cell scraper and transferred to a 1.5ml centrifuge tube. After lysing the cells on ice for 30 minutes and ensuring complete cell lysis through pipetting, the lysate was centrifuged at 12,000 rpm for 10 minutes at 4°C, and the resulting supernatant was collected as the total protein solution. The protein concentration was determined using a microspectrophotometer, and for accurate measurement, a non-denatured protein solution was utilized with the BCA Protein Assay Kit. The protein samples were denatured by combining them with a 4:1 ratio of reducing protein loading buffer (Solarbio, P1015) and heating in a boiling water bath for 15 minutes, followed by storage at -20°C. SDS-PAGE electrophoresis was performed using a 10% polyacrylamide gel, with approximately 25μg of protein loaded per well. The gel was run at 80V for gel concentration and stacking, and then at 120V for protein separation. Following electrophoresis, the proteins were transferred onto a PVDF membrane using semi-dry transfer. The membrane was subjected to immunoreaction by incubating it in TBST with blocking solution, followed by incubation with primary and secondary antibodies. Finally, chemiluminescent detection using an ECL reagent was performed, and the resulting protein bands were analyzed using ImageJ software to determine their grayscale intensity.

#### 17. Transwell cell migration assay

Mouse pulmonary arterial smooth muscle cells were cultured in the co-culture system until they reached 70-80% confluency. The cells were then suspended in serum-free medium at an appropriate density. A polycarbonate or collagen-coated membrane filter was placed on top of the Transwell insert, which was subsequently positioned in a 24-well plate filled with culture medium. The cell suspension was added to each Transwell insert at a typical density of 1-2×10^5 cells, while separate wells containing only culture medium served as control. The Transwell plate was incubated in a cell culture incubator at 37°C with 5% CO2 for the specified incubation period. After incubation, the Transwell plate was carefully removed, and cells on the upper side of the membrane were gently wiped off. The Transwell inserts were then transferred to a new plate filled with PBS, and any excess PBS was removed by centrifugation. The cells within the Transwell inserts were fixed using a 4% paraformaldehyde or formaldehyde solution, followed by staining with a suitable staining solution such as crystal violet. The migrated cells were observed and imaged using a microscope. Quantitative analysis of cell migration was performed using ImageJ software.

#### 18. Protein microarray of inflammatory cytokines and chemokines

One hundred microliters of cell culture supernatant from each time-point and group were utilized for the detection and quantification of cytokines and chemokines. This was achieved through the utilization of a Quantibody Mouse Inflammation Array I kit (RayBiotech, Inc. Norcross, GA; Cat. No. QAM-INF-1) following the manufacturer's instructions. The signals were visualized as green fluorescence with a Cy3 wavelength of 532 nm using an InnoScan 300 Microarray Scanner (Innopsys, France). Subsequently, quantitative data analysis was performed using the RayBiotech mouse Inflammation Array 1 software (QAM-INF-1-Q-Analyzer).

**Supplementary Table E1. Clinical characteristics of the recipients included in the study.**

| **Variables** | **Recipient 1** | **Recipient 2** |
|---|---|---|
| Age, years | 56 | 53 |
| Gender | F | F |
| WHO FC | I | I |
| Onset to diagnosis, months | 12 | 3 |
| N-terminal fragmental of pro-brian natriuretic peptide, pg/ml | 7068 | 2645 |
| Blood pressure, mmHg | 110/82 | 105/60 |
| Heart rate, bpm | 85 | 87 |
| **Echocardiography** | | |
| Left ventricular end-systolic diamter, cm | 3.5 | 2.7 |
| Left ventricular end-diastolic diamter, cm | 4.1 | 4.1 |
| Left ventricular ejection fraction, % | 32 | 64 |
| Pulmonary arterial systolic pressure, mmHg | 100 | 132 |
| Pulmonary artery diameter (left), mm | - | 18 |
| Pulmonary artery diameter (right), mm | - | 19 |
| Pulmonary artery diameter (root), mm | 39 | 39 |
| Interventricular septal thickness, mm | 9 | 8 |
| Left atrium, cm | 2.9 | 3.0 |
| **Laboratory examination** | | |
| Hemoglobin, g/L | 158 | 93 |
| White blood cells, $10^9$/L | 5.6 | 4.61 |
| Lymphocyte, % | 34.1 | 10.1 |
| Neutrophils, % | 53.5 | 83.8 |
| Monocyte, % | 10.8 | 5.3 |
| Eosinophil, % | 1.1 | 0.4 |

| | | |
|---|---|---|
| Basophil, % | 0.5 | 0.4 |
| Platelet, $10^9$/L | 214 | 44 |
| Alanine aminotransferase, IU/L | 78 | 10 |
| Glutamic oxalacetic transaminase, IU/L | 82 | 17 |
| Total Billirubin, µmol/L | 22.3 | 50.9 |
| Direct Billirubin, µmol/L | 12.1 | 27.6 |
| Creatinine, µmol/L | 104 | 66 |
| Uric Acic, µmol/L | 373 | 570 |
| D-Dimer, ng/mL | 2786 | 270 |

**Supplementary Table E2. Clinical characteristics of the donors included in the study.**

| Variables | Donor 1 | Donor 2 | Donor 3 |
|---|---|---|---|
| Age, years | 45 | 21 | 49 |
| Gender | M | M | M |
| BMI, kg/㎡ | 23.5 | 20.8 | 18.4 |
| Cause of death | Brain Injury | Brain Injury | Stroke |
| **past medical history** | | | |
| Former smoker (True/False) | T | F | F |
| Primary pulmonary disease (True/False) | F | F | F |
| Hypertension (True/False) | F | F | F |
| Coronary artery disease (True/False) | F | F | F |
| Diabetes mellitus (True/False) | F | F | F |
| Malignant tumor (True/False) | F | F | F |
| HIV (True/False) | F | F | F |
| **Laboratory examination** | | | |
| CMV-IgG (Negative/Positive) | P | N | P |

| | | | |
|---|---|---|---|
| CMV-IgM (Negative/Positive) | N | N | N |
| Hepatitis B surface antigen (Negative/Positive) | N | N | N |
| HCV-DNA (Negative/Positive) | N | N | N |
| EBV-IgG (Negative/Positive) | P | N | P |
| EBV-IgM (Negative/Positive) | N | N | N |
| Sputum smear for tuberculosis (Negative/Positive) | N | N | N |
| Xpert test for tuberculosis (Negative/Positive) | N | N | N |
| Blood culture (Negative/Positive) | N | N | N |
| White blood cells, $10^9$/L | 11.76 | 23.7 | 31.5 |
| C-reactive protein, mg/L | 276 | 43.3 | 33.6 |
| Procalcitonin, ug/L | 15.8 | 7.7 | 5.3 |

HIV, human immunodeficiency virus; CMV, cytomegalovirus; HCV, hepatitis C virus; EBV, Epstein-Barr virus; Xpert, Xpert MTB/RIF（Mycobacterium tuberculosis/Rifampicin）.

**Supplementary Table E3. Evaluation of experimental animal models of pulmonary arterial hypertension.**

By using cardiac ultrasound examination, evaluate the physiological indicators of the control group and the model group mice.

**Supplementary Table E4. Markers (by avg_logFC) for each cluster of fibroblasts and myofibrocytes in normal pulmonary artery.**

Differentially expressed genes (DEGs) of subgroups from clustering fibroblasts and myofibrocytes in normal pulmonary arteries. The DEGs reveal distinct subpopulation patterns.

**Supplementary Table E5. Markers (by avg_logFC) for each cluster of macrophages in normal pulmonary artery.**

Differentially expressed genes of subgroups from clustering macrophages in normal pulmonary arteries. The DEGs reveal distinct subpopulation patterns.

**Supplementary Table E6. Markers (by avg_logFC) for each cluster of endothelial cells in normal pulmonary artery**

Differentially expressed genes of normal and (endothelial-to-mesenchymal transition) EndMT endothelial cells in normal pulmonary arteries. The DEGs reveal distinct subpopulation patterns.

**Supplementary Table E7. Markers (by avg_logFC) for each cluster of smooth muscle cells in normal pulmonary artery**

Differentially expressed genes of clusters of smooth muscle cells with different phenotypes (contractile, chondroid-like, and fibroblast-like) in normal pulmonary arteries. The DEGs reveal distinct subpopulation patterns.

**Supplementary Table E8. Markers (by avg_logFC) for each cluster of smooth muscle cells in normal and hypertensive pulmonary artery**

Differentially expressed genes of clusters of smooth muscle cells with different phenotypes (contractile, chondroid-like, and fibroblast-like) in the integration data of normal and hypertensive pulmonary arteries. The DEGs reveal distinct subpopulation patterns.

**References**

1. Jiang, L. et al. Nonbone Marrow CD34+ Cells Are Crucial for Endothelial Repair of Injured Artery. Circulation Research **129**, e146–e165 (2021).
2. Stuart, T. et al. Comprehensive Integration of Single-Cell Data. Cell **177**, 1888-1902.e21 (2019).
3. Germain, P.-L., Lun, A., Garcia Meixide, C., Macnair, W. & Robinson, M. D. Doublet identification in single-cell sequencing data using scDblFinder. F1000Res **10**, 979 (2021).
4. Wu, T. et al. clusterProfiler 4.0: A universal enrichment tool for interpreting omics data. Innovation (Camb) **2**, 100141 (2021).
5. Yu, G., Wang, L.-G., Han, Y. & He, Q.-Y. clusterProfiler: an R package for comparing

biological themes among gene clusters. OMICS **16**, 284–287 (2012).

6. Holland, C. H., Szalai, B. & Saez-Rodriguez, J. Transfer of regulatory knowledge from human to mouse for functional genomics analysis. Biochim Biophys Acta Gene Regul Mech **1863**, 194431 (2020).

7. Trapnell, C. et al. The dynamics and regulators of cell fate decisions are revealed by pseudotemporal ordering of single cells. Nat Biotechnol **32**, 381–386 (2014).

8. X, Q. et al. Single-cell mRNA quantification and differential analysis with Census. Nature methods **14**, (2017).

9. Garcia-Alonso, L., Holland, C. H., Ibrahim, M. M., Turei, D. & Saez-Rodriguez, J. Benchmark and integration of resources for the estimation of human transcription factor activities. Genome Res **29**, 1363–1375 (2019).

10. Holland, C. H. et al. Robustness and applicability of transcription factor and pathway analysis tools on single-cell RNA-seq data. Genome Biol **21**, 36 (2020).

11. Alvarez, M. J. et al. Functional characterization of somatic mutations in cancer using network-based inference of protein activity. Nat Genet **48**, 838–847 (2016).

12. Jin, S. et al. Inference and analysis of cell-cell communication using CellChat. Nat Commun **12**, 1088 (2021).

**Supplementary figure legends:**

**Figure E1: Transcriptomic characteristics of fibroblasts and macrophages.** (A) Correlation assessment of fibroblast and myofibrocyte subpopulations. (B) Dot plot visualization of functional enrichment analysis using KEGG and Gene Ontology (GO) terms. Enriched KEGG pathways and GO terms were identified based on differentially expressed genes (DEGs) of fibroblast and myofibrocyte subpopulations. (C) Marker genes used for cell definition, including classical markers for fibroblasts and myofibrocytes. (D) Scatter plot illustrating the differentially expressed genes between M1 macrophage and M2 macrophage.

**Figure E2: Identification of endothelial cell subpopulation-specific genes and dynamic gene expression during endothelial-to-mesenchymal transition.** (A) Dot plot showing the top 5 genes with the highest average expression and percentage of expressing cells for each endothelial cell subpopulation. (B) Heatmap displaying the expression of genes that vary along the pseudo-temporal trajectory of endothelial-to-mesenchymal transition, which were grouped into 3 clusters based on their distinct expression profiles.

**Figure E3: Endothelial cells from various tissues.** (A) UMAP plot of endothelial cells based on their gene expression profiles. Cells from different clusters are colored differently. (B) Feature plots of selected genes that distinguish normal endothelial cells (normal ECs, first rows) and endothelial cells undergoing endothelial-to-mesenchymal transition (EndMT ECs, second and third rows). The gene expression levels are represented by the color intensity of the dots. (C) Correlation matrix of endothelial cell clusters. (D) Violin plot showing the expression distribution of canonical markers for endothelial cells and mesenchymal cells. (E) Dot plot displaying the top 5 genes with the highest average expression and percentage of expressing cells for each cluster.

**Figure E4: Functional enrichment analysis of endothelial cells from various tissues.** Dot plot showing the top enriched GO and KEGG terms for normal ECs (A) and EndMT ECs (B) originating from different tissues.

**Figure E5: Comparative gene expression profiles of smooth muscle cells with distinct phenotypes.** (A) Dot plot showing the top 10 genes with the highest average expression and percentage of expressing cells for each smooth muscle cell cluster. (B) Scatter plot illustrating the differentially expressed genes between contractile SMC and chondroid-like SMC (upper panel), as well as contractile SMC and fibroblast-like SMC (lower panel).

**Figure E6: Smooth muscle cells from various tissues.** (A) UMAP dimensionality reduction analysis of smooth muscle cells based on their gene expression profiles. Cells from different clusters are shown in different colors. (B) Feature plots of some genes that can distinguish contractile SMC (CNN1, TAGLN, MYH11, and ACTA2), fibroblast-like SMC (DCN and GSN), and chondroid-like SMC (LGALS3 and BMP2). The gene expression levels are indicated by the color intensity of the dots. (C) Functional enrichment analysis using KEGG and Gene Ontology (GO) terms, and visualization with dot plots. Enriched KEGG pathways and GO terms were determined based on differentially expressed genes (DEGs) of contractile SMC (SMC), chondroid-like SMC (C-SMC), and fibroblast-like SMC (F-SMC).

**Figure E7: Gene expression profiles of smooth muscle cell subtypes and phenotypic transition in normal and hypertensive groups.** (A) Violin plot depicting the top 10 differentially expressed genes in each smooth muscle cell subtype between the control and hypertensive groups. (B) Expression patterns of selected genes along the pseudotime trajectory in the control and hypertensive groups. (C) Heatmap illustrating the expression levels of genes that exhibit dynamic changes along the pseudotemporal trajectory of smooth muscle cell phenotypic switching, classified into three distinct clusters based on their expression profiles.

**Figure E8: Cell–cell interactions and macrophage markers in normal and hypertensive groups.** (A) Scatter plot of the outgoing and incoming interaction intensity in two-dimensional space. The circle size indicates the number of significant receptor-ligand pathways for each cell type. (B) Circle plots of the interaction number and strength of VISFATIN (NAMPT) signaling pathway between normal and hypertensive groups. (C) Heatmap of the relative strength of incoming and outgoing signal pathway networks for each cluster. The vertical axis shows the source or target cell, and the horizontal axis shows the pathway that sends or receives the signal. The heatmap color represents the signal strength. The bars on the top and right sides show the sum of the signal strength for each cell or pathway. (D) Feature plot of the expression of canonical markers for macrophage subtypes.

**Figure E9: NAMPT expression of macrophages and the secretion profile with siNAMPT treatment.** (A) Immunofluorescence staining of normal and sugen5416+hypoxia mouse arteries (scale bar, 100 μm). Co-stained cells are indicated by blue arrows. (B) A dot plot showing the top enriched GO terms in biological process (BP), cellular component (CC) and molecular function (MF) categories based on the differential expression of chemokines and cytokines secreted by macrophages in the co-culture system with siNAMPT or siNC, as measured by protein array analysis. The dot size reflects the number of genes in each pathway, and the dot color denotes the adjusted p-value, ranging from low (green) to high (red).

**Figure E10: Effect of siNAMPT/siNC-treated macrophages on smooth muscle cells.** (A) Immunofluorescence staining of SMCs in the siNC and siNAMPT groups, with a scale bar of 100 μm. (B) Western blot analysis of P65 and P-p65 protein expression in smooth muscle cells.

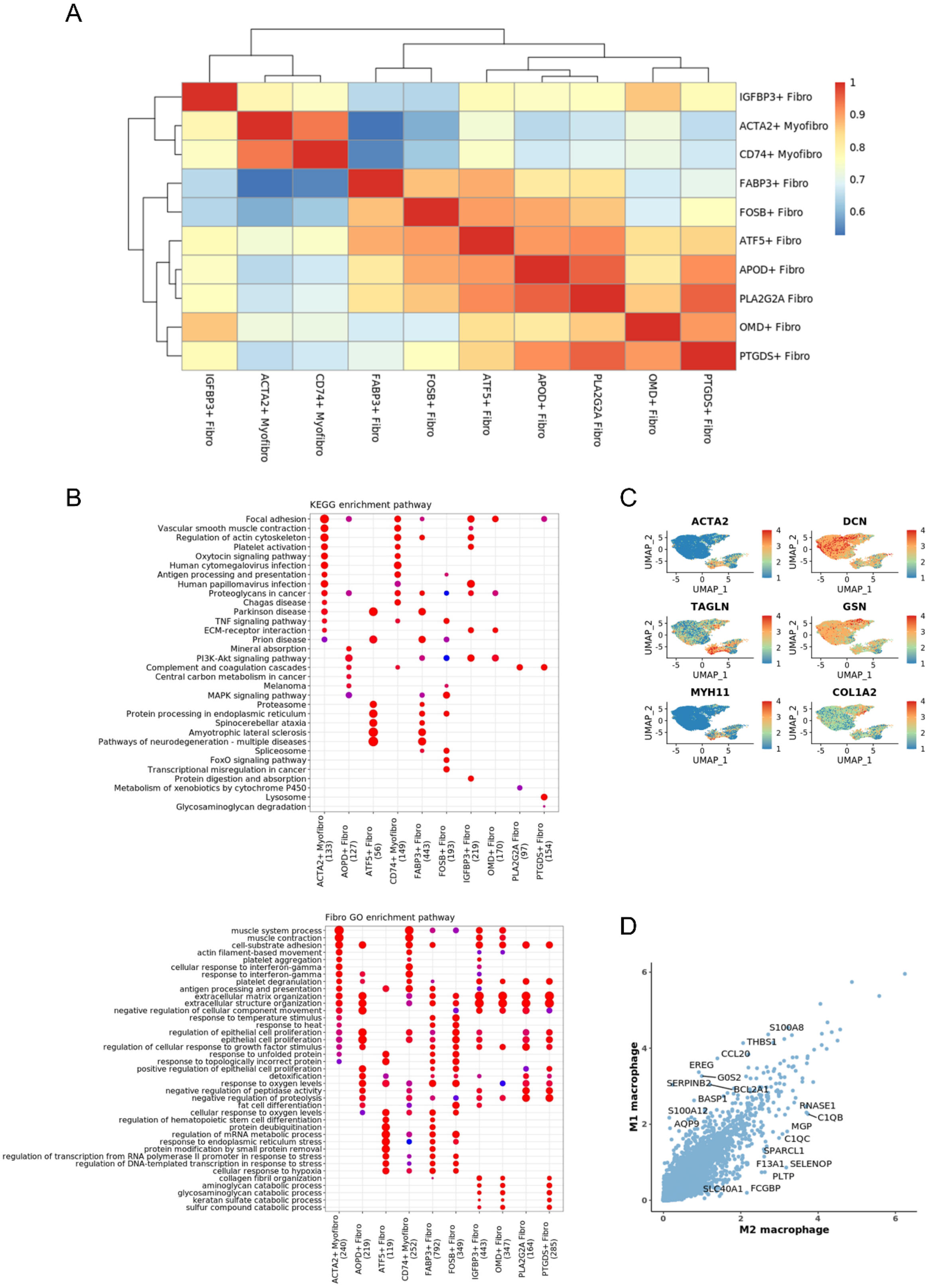
A
IGFBP3+ Fibro
ACTA2+ Myofibro
CD74+ Myofibro
FABP3+ Fibro
FOSB+ Fibro
ATF5+ Fibro
APOD+ Fibro
PLA2G2A Fibro
OMD+ Fibro
PTGDS+ Fibro
B
KEGG enrichment pathway
Fibro GO enrichment pathway
C
ACTA2
DCN
TAGLN
GSN
MYH11
COL1A2
UMAP_1
UMAP_2
D
M1 macrophage
M2 macrophage

**Figure E1**

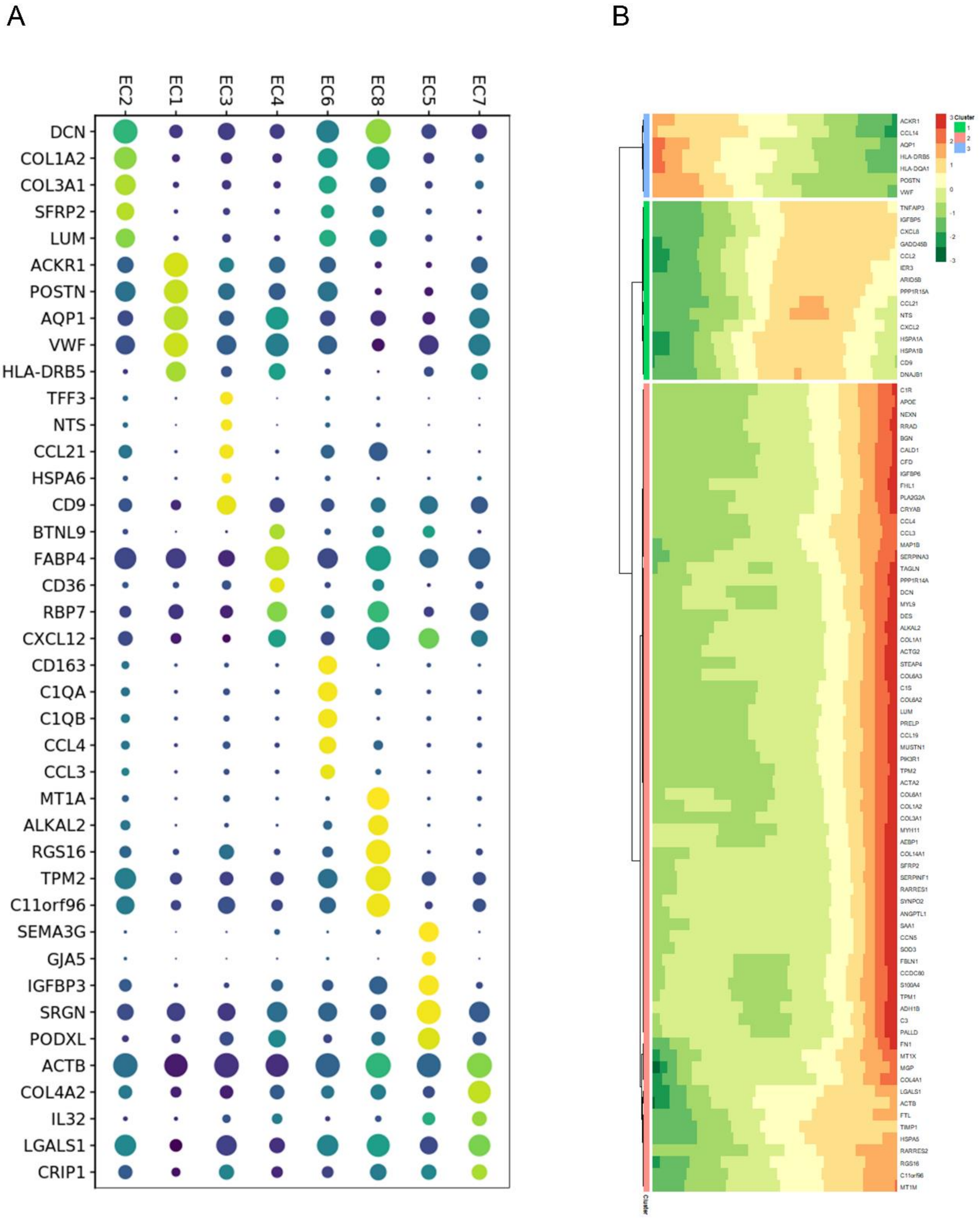
A
B
EC2
EC1
EC3
EC4
EC6
EC8
EC5
EC7
DCN
COL1A2
COL3A1
SFRP2
LUM
ACKR1
POSTN
AQP1
VWF
HLA-DRB5
TFF3
NTS
CCL21
HSPA6
CD9
BTNL9
FABP4
CD36
RBP7
CXCL12
CD163
C1QA
C1QB
CCL4
CCL3
MT1A
ALKAL2
RGS16
TPM2
C11orf96
SEMA3G
GJA5
IGFBP3
SRGN
PODXL
ACTB
COL4A2
IL32
LGALS1
CRIP1
ACKR1
CCL14
AQP1
HLA-DRB5
HLA-DQA1
POSTN
VWF
TNFAIP3
IGFBP5
CXCL8
GADD45B
CCL2
IER3
ARID5B
PPP1R15A
CCL21
NTS
CXCL2
HSPA1A
HSPA1B
CD9
DNAJB1
C1R
APOE
NEXN
RRAD
BGN
CALD1
CFD
IGFBP6
FHL1
PLA2G2A
CRYAB
CCL4
CCL3
MAP1B
SERPINA3
TAGLN
PPP1R14A
DCN
MYL9
DES
ALKAL2
COL1A1
ACTG2
STEAP4
COL6A3
C1S
COL6A2
LUM
PRELP
CCL19
MUSTN1
PIK3R1
TPM2
ACTA2
COL6A1
COL1A2
COL3A1
MYH11
AEBP1
COL14A1
SFRP2
SERPINF1
RARRES1
SYNPO2
ANGPTL1
SAA1
CCN5
SOD3
FBLN1
CCDC80
S100A4
TPM1
ADH1B
C3
PALLD
FN1
MT1X
MGP
COL4A1
LGALS1
ACTB
FTL
TIMP1
HSPA5
RARRES2
RGS16
C11orf96
MT1M
Cluster
3
2
1
0
-1
-2
-3
Cluster
1
2
3
Cluster

**Figure E2**

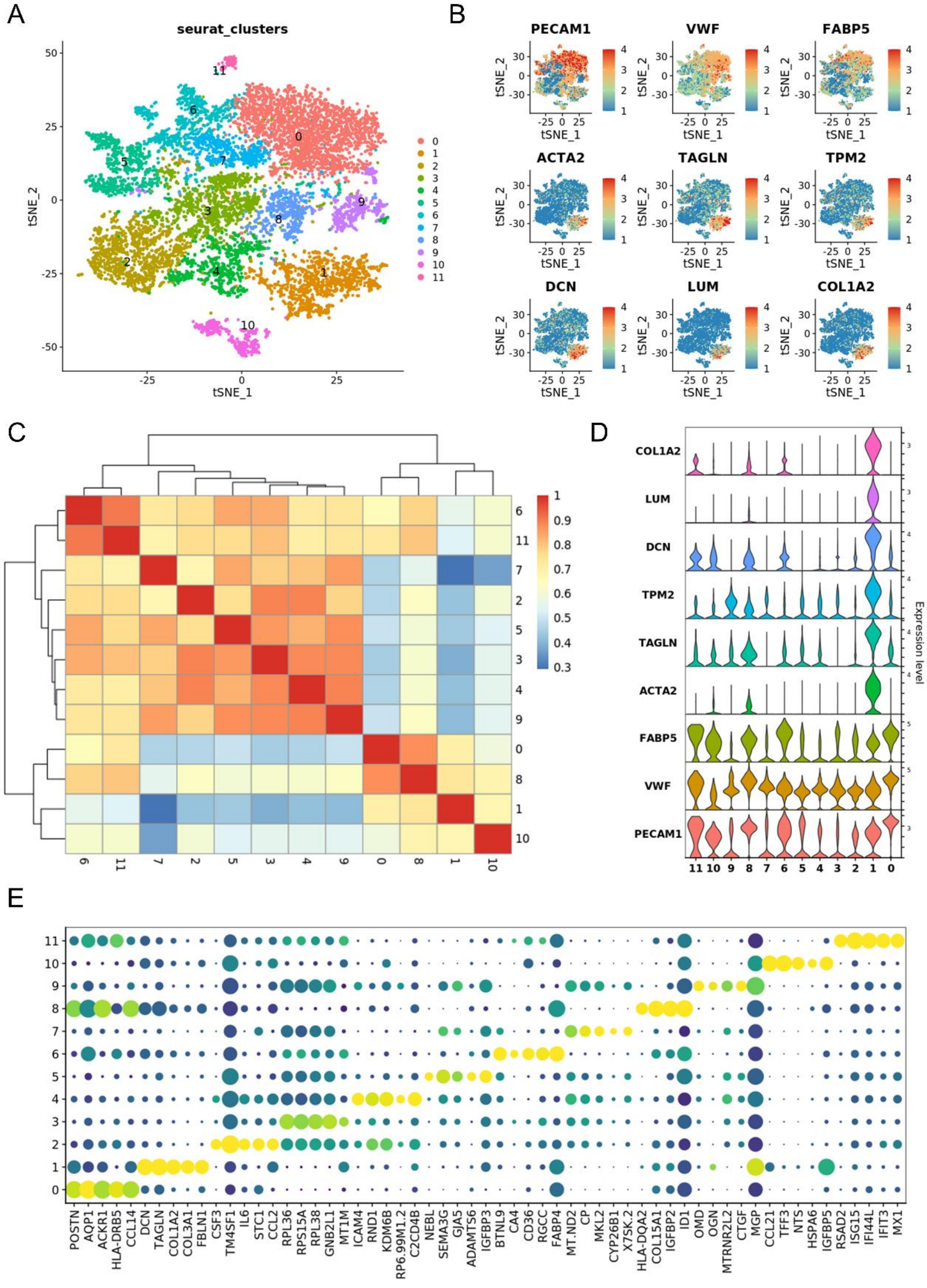
A
seurat_clusters
tSNE_1
tSNE_2
B
PECAM1
VWF
FABP5
ACTA2
TAGLN
TPM2
DCN
LUM
COL1A2
C
D
COL1A2
LUM
DCN
TPM2
TAGLN
ACTA2
FABP5
VWF
PECAM1
Expression level
E
POSTN
AQP1
ACKR1
HLA-DRB5
CCL14
DCN
TAGLN
COL1A2
COL3A1
FBLN1
CSF3
TM4SF1
IL6
STC1
CCL2
RPL36
RPS15A
RPL38
GNB2L1
MT1M
ICAM4
RND1
KDM6B
RP6.99M1.2
C2CD4B
NEBL
SEMA3G
GJA5
ADAMTS6
IGFBP3
BTNL9
CA4
CD36
RGCC
FABP4
MT.ND2
CP
MKL2
CYP26B1
X7SK.2
HLA-DQA2
COL15A1
IGFBP2
ID1
OMD
OGN
MTRNR2L2
CTGF
MGP
CCL21
TFF3
NTS
HSPA6
IGFBP5
RSAD2
ISG15
IFI44L
IFIT3
MX1

**Figure E3**

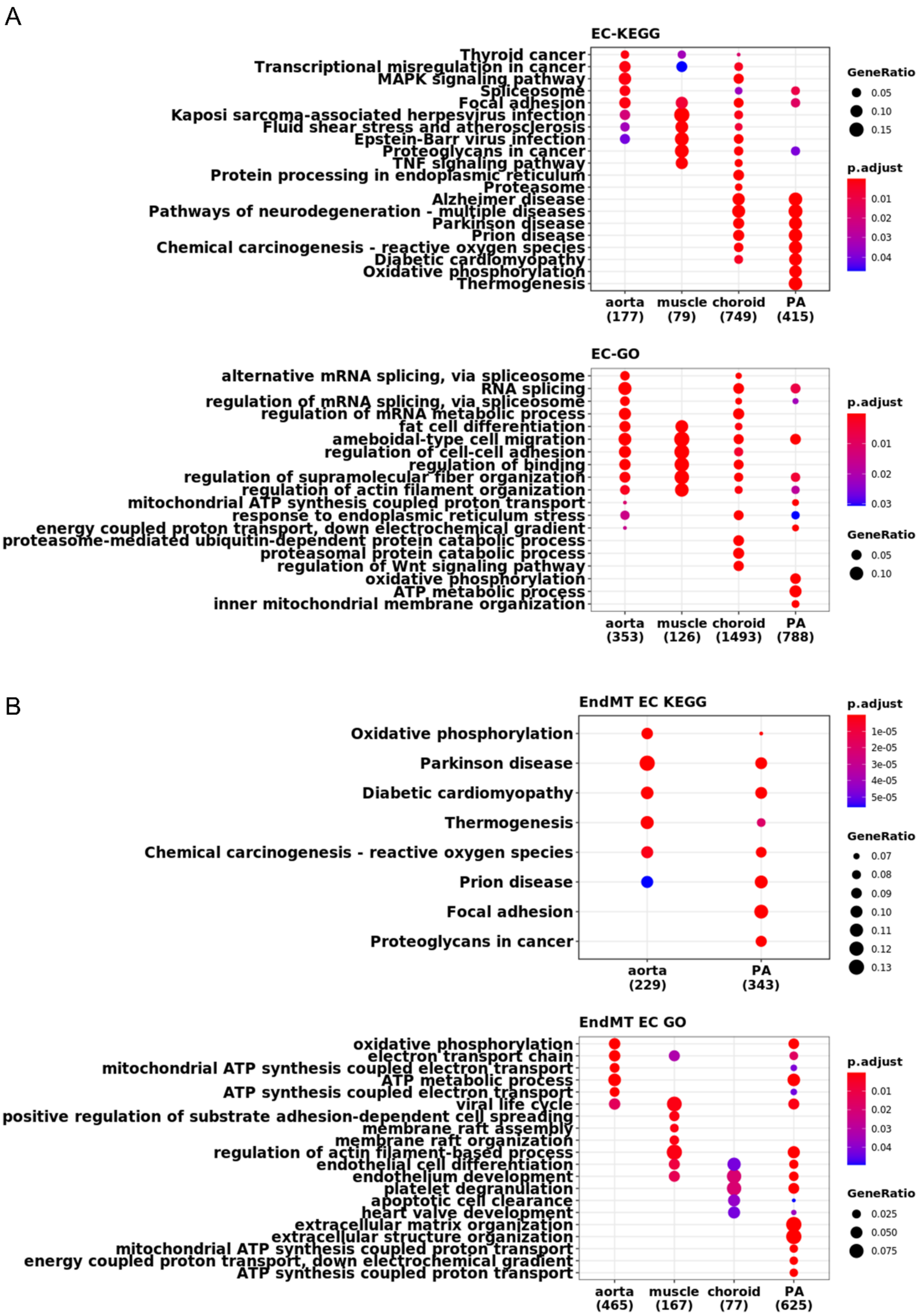
A
EC-KEGG
Thyroid cancer
Transcriptional misregulation in cancer
MAPK signaling pathway
Spliceosome
Focal adhesion
Kaposi sarcoma-associated herpesvirus infection
Fluid shear stress and atherosclerosis
Epstein-Barr virus infection
Proteoglycans in cancer
TNF signaling pathway
Protein processing in endoplasmic reticulum
Proteasome
Alzheimer disease
Pathways of neurodegeneration - multiple diseases
Parkinson disease
Prion disease
Chemical carcinogenesis - reactive oxygen species
Diabetic cardiomyopathy
Oxidative phosphorylation
Thermogenesis
aorta (177)
muscle (79)
choroid (749)
PA (415)
GeneRatio
0.05
0.10
0.15
p.adjust
0.01
0.02
0.03
0.04
EC-GO
alternative mRNA splicing, via spliceosome
RNA splicing
regulation of mRNA splicing, via spliceosome
regulation of mRNA metabolic process
fat cell differentiation
ameboidal-type cell migration
regulation of cell-cell adhesion
regulation of binding
regulation of supramolecular fiber organization
regulation of actin filament organization
mitochondrial ATP synthesis coupled proton transport
response to endoplasmic reticulum stress
energy coupled proton transport, down electrochemical gradient
proteasome-mediated ubiquitin-dependent protein catabolic process
proteasomal protein catabolic process
regulation of Wnt signaling pathway
oxidative phosphorylation
ATP metabolic process
inner mitochondrial membrane organization
aorta (353)
muscle (126)
choroid (1493)
PA (788)
p.adjust
0.01
0.02
0.03
GeneRatio
0.05
0.10
B
EndMT EC KEGG
Oxidative phosphorylation
Parkinson disease
Diabetic cardiomyopathy
Thermogenesis
Chemical carcinogenesis - reactive oxygen species
Prion disease
Focal adhesion
Proteoglycans in cancer
aorta (229)
PA (343)
p.adjust
1e-05
2e-05
3e-05
4e-05
5e-05
GeneRatio
0.07
0.08
0.09
0.10
0.11
0.12
0.13
EndMT EC GO
oxidative phosphorylation
electron transport chain
mitochondrial ATP synthesis coupled electron transport
ATP metabolic process
ATP synthesis coupled electron transport
viral life cycle
positive regulation of substrate adhesion-dependent cell spreading
membrane raft assembly
membrane raft organization
regulation of actin filament-based process
endothelial cell differentiation
endothelium development
platelet degranulation
apoptotic cell clearance
heart valve development
extracellular matrix organization
extracellular structure organization
mitochondrial ATP synthesis coupled proton transport
energy coupled proton transport, down electrochemical gradient
ATP synthesis coupled proton transport
aorta (465)
muscle (167)
choroid (77)
PA (625)
p.adjust
0.01
0.02
0.03
0.04
GeneRatio
0.025
0.050
0.075

**Figure E4**

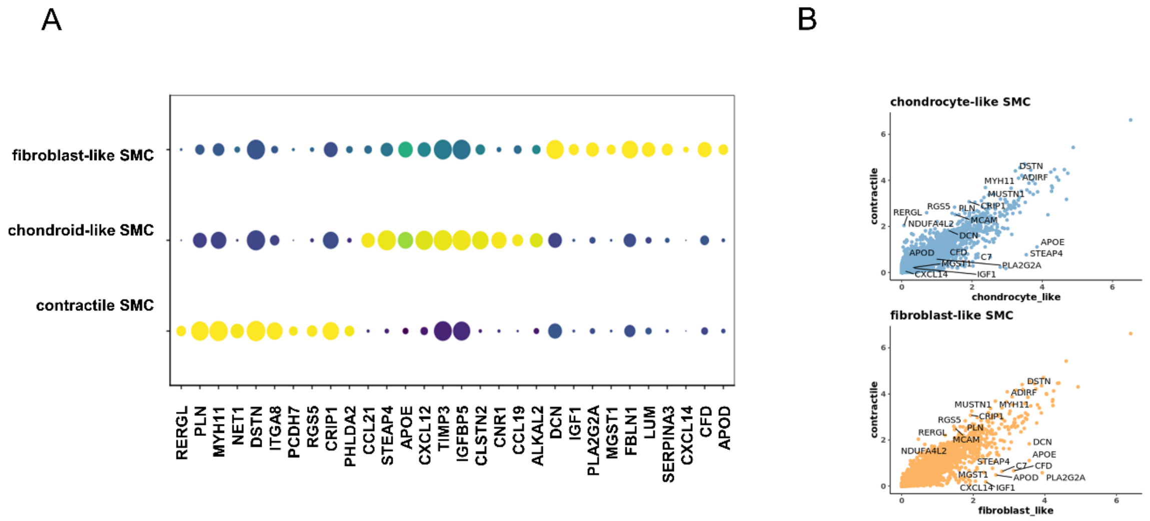

A
B
fibroblast-like SMC
chondroid-like SMC
contractile SMC
RERGL
PLN
MYH11
NET1
DSTN
ITGA8
PCDH7
RGS5
CRIP1
PHLDA2
CCL21
STEAP4
APOE
CXCL12
TIMP3
IGFBP5
CLSTN2
CNR1
CCL19
ALKAL2
DCN
IGF1
PLA2G2A
MGST1
FBLN1
LUM
SERPINA3
CXCL14
CFD
APOD
chondrocyte-like SMC
contractile
chondrocyte_like
fibroblast-like SMC
contractile
fibroblast_like

**Figure E5**

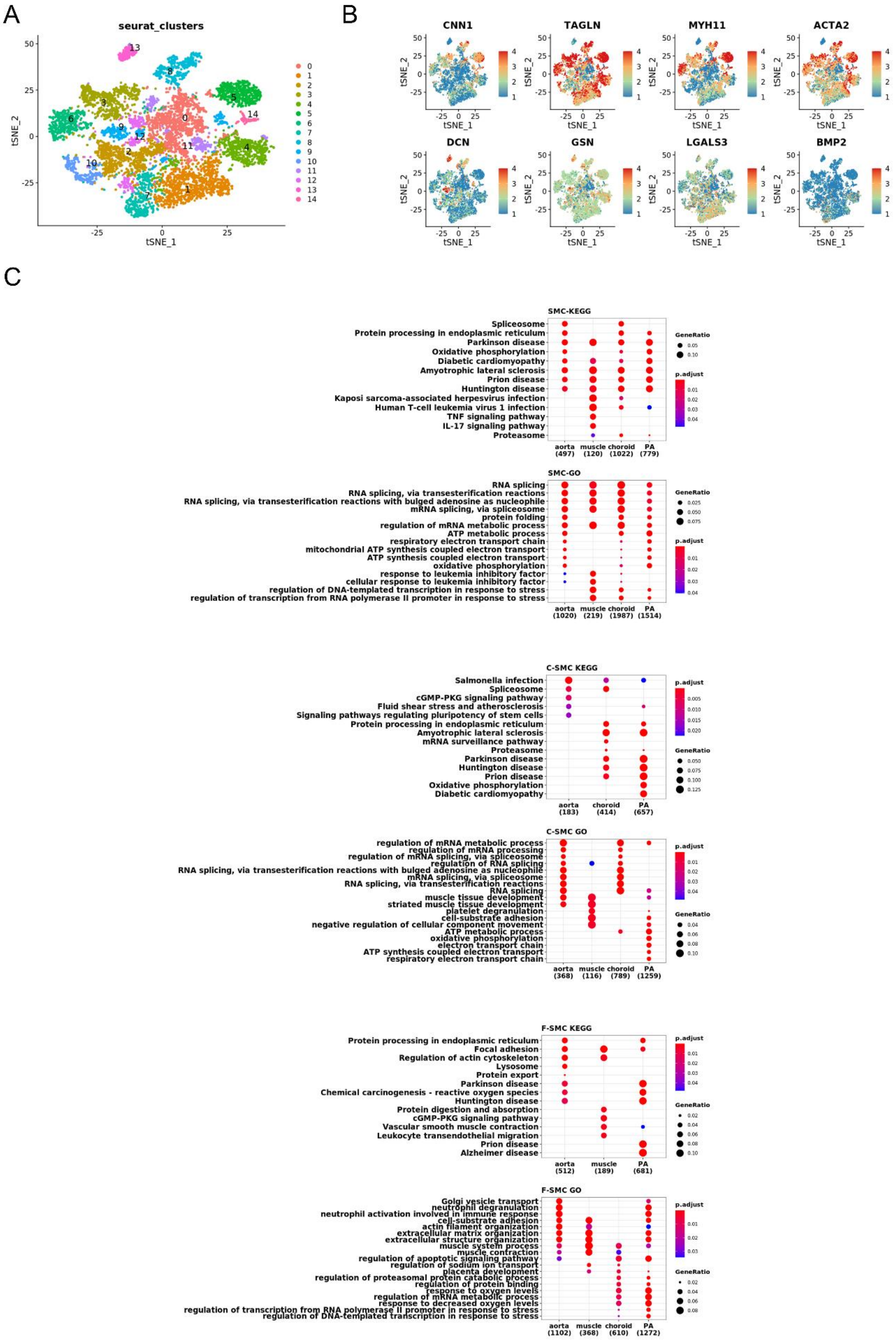
A
seurat_clusters
tSNE_1
tSNE_2
B
CNN1
TAGLN
MYH11
ACTA2
DCN
GSN
LGALS3
BMP2
C
SMC-KEGG
Spliceosome
Protein processing in endoplasmic reticulum
Parkinson disease
Oxidative phosphorylation
Diabetic cardiomyopathy
Amyotrophic lateral sclerosis
Prion disease
Huntington disease
Kaposi sarcoma-associated herpesvirus infection
Human T-cell leukemia virus 1 infection
TNF signaling pathway
IL-17 signaling pathway
Proteasome
aorta (497)
muscle (120)
choroid (1022)
PA (779)
GeneRatio
p.adjust
SMC-GO
RNA splicing
RNA splicing, via transesterification reactions
RNA splicing, via transesterification reactions with bulged adenosine as nucleophile
mRNA splicing, via spliceosome
protein folding
regulation of mRNA metabolic process
ATP metabolic process
respiratory electron transport chain
mitochondrial ATP synthesis coupled electron transport
ATP synthesis coupled electron transport
oxidative phosphorylation
response to leukemia inhibitory factor
cellular response to leukemia inhibitory factor
regulation of DNA-templated transcription in response to stress
regulation of transcription from RNA polymerase II promoter in response to stress
aorta (1020)
muscle (219)
choroid (1987)
PA (1514)
C-SMC KEGG
Salmonella infection
Spliceosome
cGMP-PKG signaling pathway
Fluid shear stress and atherosclerosis
Signaling pathways regulating pluripotency of stem cells
Protein processing in endoplasmic reticulum
Amyotrophic lateral sclerosis
mRNA surveillance pathway
Proteasome
Parkinson disease
Huntington disease
Prion disease
Oxidative phosphorylation
Diabetic cardiomyopathy
aorta (183)
choroid (414)
PA (657)
C-SMC GO
regulation of mRNA metabolic process
regulation of mRNA processing
regulation of mRNA splicing, via spliceosome
regulation of RNA splicing
RNA splicing, via transesterification reactions with bulged adenosine as nucleophile
mRNA splicing, via spliceosome
RNA splicing, via transesterification reactions
RNA splicing
muscle tissue development
striated muscle tissue development
platelet degranulation
cell-substrate adhesion
negative regulation of cellular component movement
ATP metabolic process
oxidative phosphorylation
electron transport chain
ATP synthesis coupled electron transport
respiratory electron transport chain
aorta (368)
muscle (116)
choroid (789)
PA (1259)
F-SMC KEGG
Protein processing in endoplasmic reticulum
Focal adhesion
Regulation of actin cytoskeleton
Lysosome
Protein export
Parkinson disease
Chemical carcinogenesis - reactive oxygen species
Huntington disease
Protein digestion and absorption
cGMP-PKG signaling pathway
Vascular smooth muscle contraction
Leukocyte transendothelial migration
Prion disease
Alzheimer disease
aorta (512)
muscle (189)
PA (681)
F-SMC GO
Golgi vesicle transport
neutrophil degranulation
neutrophil activation involved in immune response
cell-substrate adhesion
actin filament organization
extracellular matrix organization
extracellular structure organization
muscle system process
muscle contraction
regulation of apoptotic signaling pathway
regulation of sodium ion transport
placenta development
regulation of proteasomal protein catabolic process
regulation of protein binding
response to oxygen levels
regulation of mRNA metabolic process
response to decreased oxygen levels
regulation of transcription from RNA polymerase II promoter in response to stress
regulation of DNA-templated transcription in response to stress
aorta (1102)
muscle (368)
choroid (610)
PA (1272)

**Figure E6**

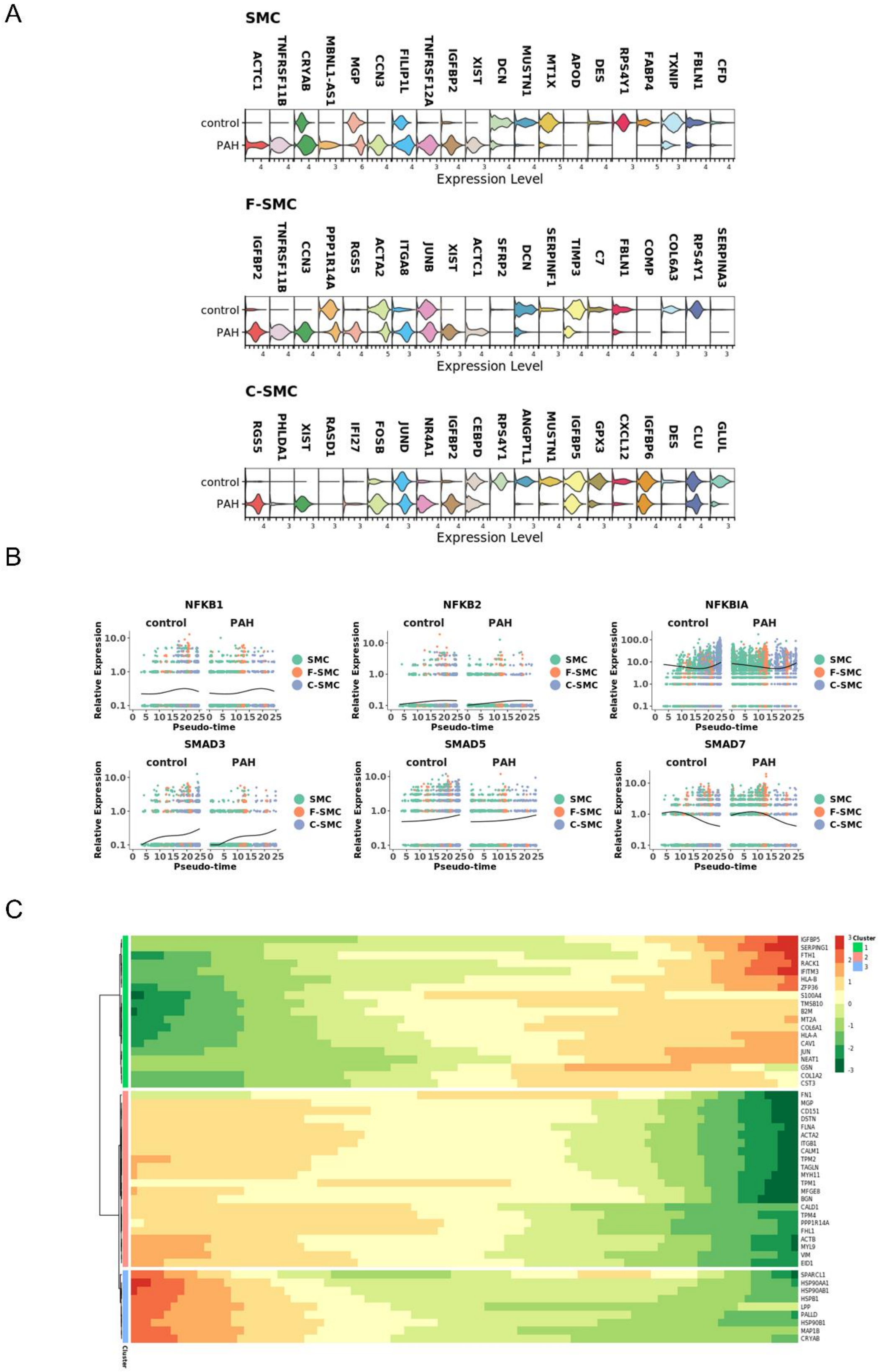
A
SMC
ACTC1
TNFRSF11B
CRYAB
MBNL1-AS1
MGP
CCN3
FILIP1L
TNFRSF12A
IGFBP2
XIST
DCN
MUSTN1
MT1X
APOD
DES
RPS4Y1
FABP4
TXNIP
FBLN1
CFD
control
PAH
Expression Level
F-SMC
IGFBP2
TNFRSF11B
CCN3
PPP1R14A
RGS5
ACTA2
ITGA8
JUNB
XIST
ACTC1
SFRP2
DCN
SERPINF1
TIMP3
C7
FBLN1
COMP
COL6A3
RPS4Y1
SERPINA3
control
PAH
Expression Level
C-SMC
RGS5
PHLDA1
XIST
RASD1
IFI27
FOSB
JUND
NR4A1
IGFBP2
CEBPD
RPS4Y1
ANGPTL1
MUSTN1
IGFBP5
GPX3
CXCL12
IGFBP6
DES
CLU
GLUL
control
PAH
Expression Level
B
NFKB1
NFKB2
NFKBIA
SMAD3
SMAD5
SMAD7
control
PAH
Relative Expression
Pseudo-time
SMC
F-SMC
C-SMC
C
IGFBP5
SERPING1
FTH1
RACK1
IFITM3
HLA-B
ZFP36
S100A4
TMSB10
B2M
MT2A
COL6A1
HLA-A
CAV1
JUN
NEAT1
GSN
COL1A2
CST3
FN1
MGP
CD151
DSTN
FLNA
ACTA2
ITGB1
CALM1
TPM2
TAGLN
MYH11
TPM1
MFGE8
BGN
CALD1
TPM4
PPP1R14A
FHL1
ACTB
MYL9
VIM
EID1
SPARCL1
HSP90AA1
HSP90AB1
HSPB1
LPP
PALLD
HSP90B1
MAP1B
CRYAB
Cluster
1
2
3

**Figure E7**

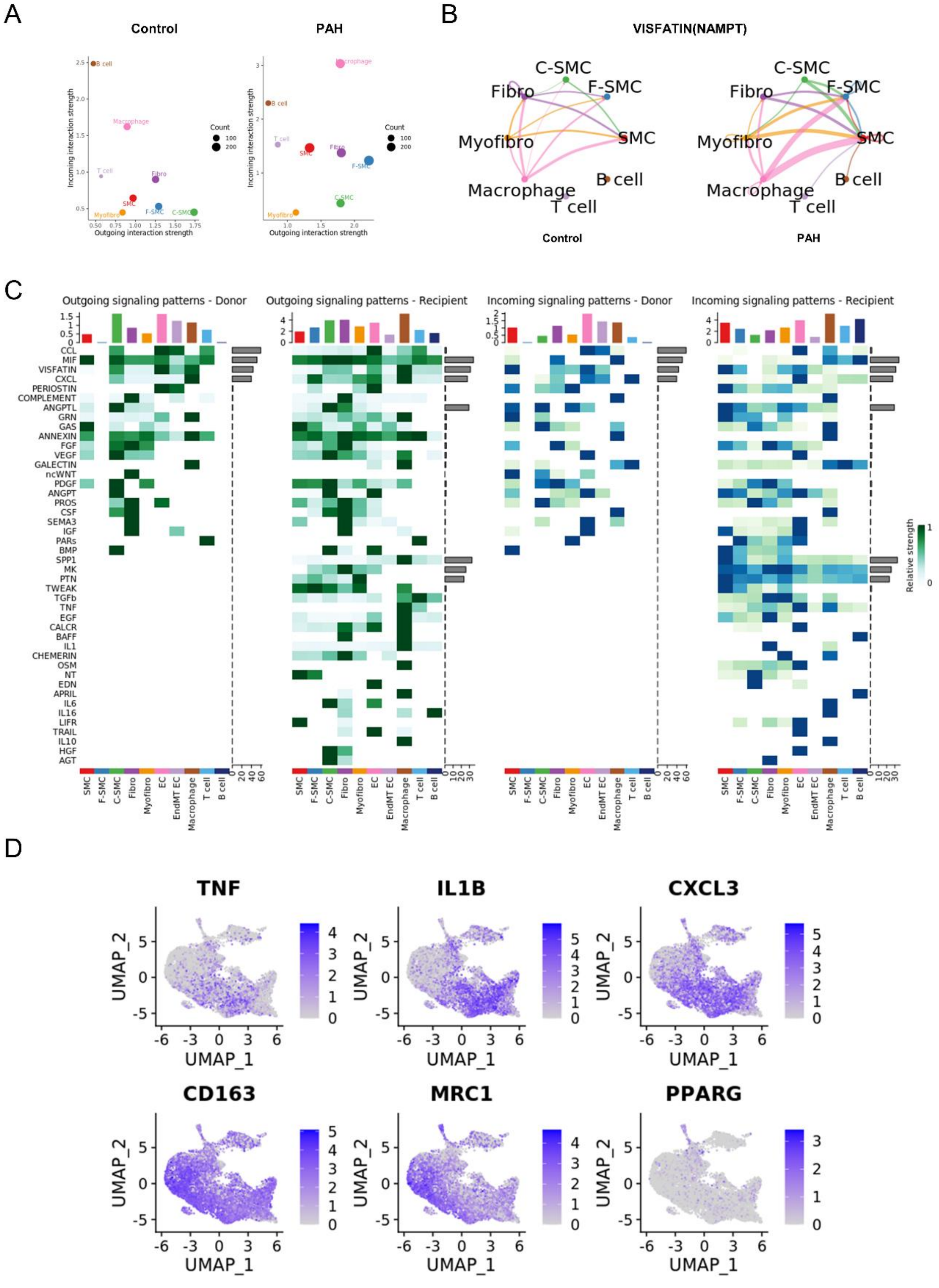
A
Control
PAH
B
VISFATIN(NAMPT)
Control
PAH
C
Outgoing signaling patterns - Donor
Outgoing signaling patterns - Recipient
Incoming signaling patterns - Donor
Incoming signaling patterns - Recipient
Relative strength
D
TNF
IL1B
CXCL3
CD163
MRC1
PPARG
UMAP_1
UMAP_2

**Figure E8**

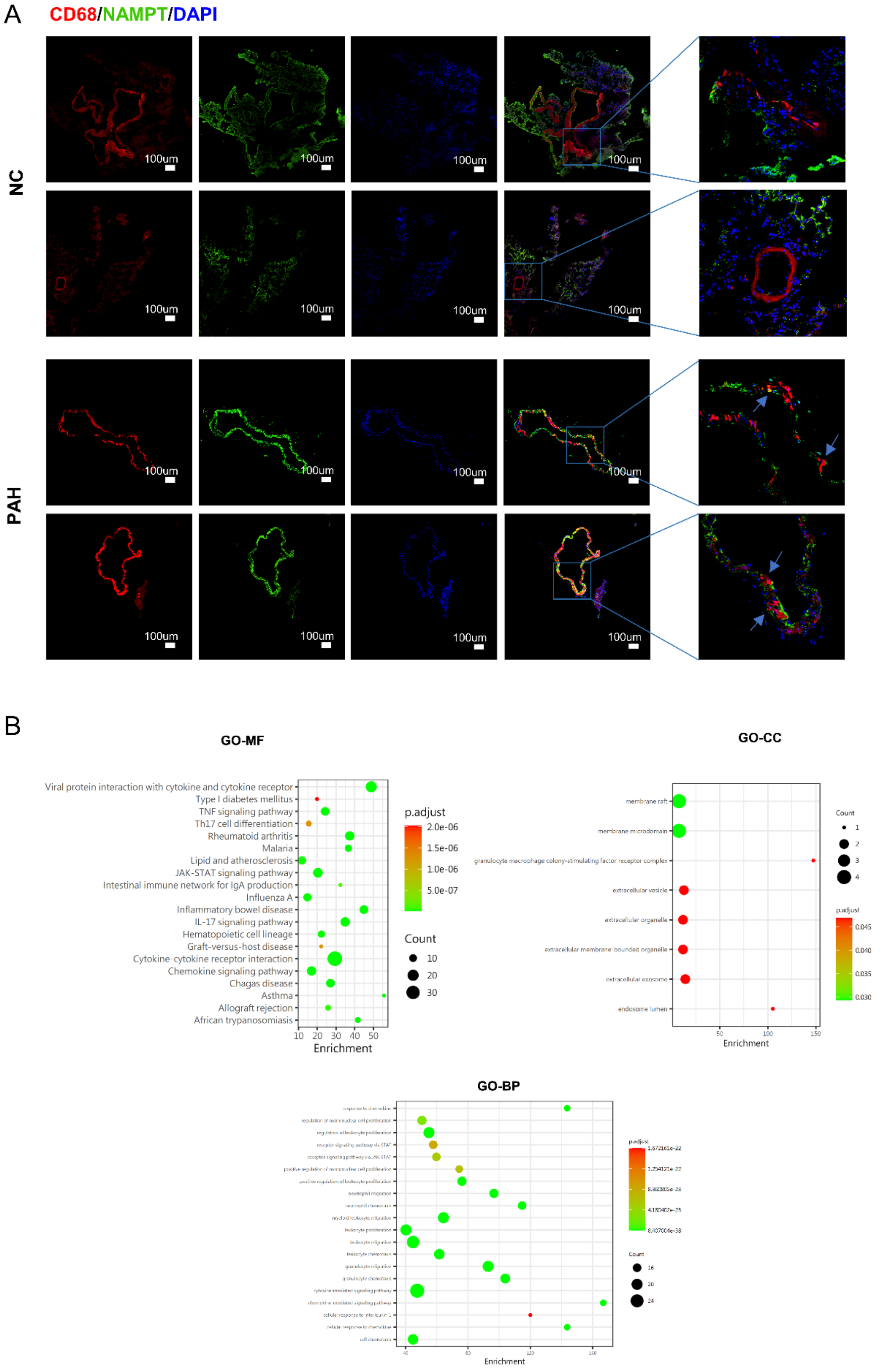
A
CD68/NAMPT/DAPI
NC
PAH
100um
B
GO-MF
Viral protein interaction with cytokine and cytokine receptor
Type I diabetes mellitus
TNF signaling pathway
Th17 cell differentiation
Rheumatoid arthritis
Malaria
Lipid and atherosclerosis
JAK-STAT signaling pathway
Intestinal immune network for IgA production
Influenza A
Inflammatory bowel disease
IL-17 signaling pathway
Hematopoietic cell lineage
Graft-versus-host disease
Cytokine-cytokine receptor interaction
Chemokine signaling pathway
Chagas disease
Asthma
Allograft rejection
African trypanosomiasis
Enrichment
p.adjust
2.0e-06
1.5e-06
1.0e-06
5.0e-07
Count
10
20
30
GO-CC
membrane raft
membrane microdomain
granulocyte macrophage colony-stimulating factor receptor complex
extracellular vesicle
extracellular organelle
extracellular membrane-bounded organelle
extracellular exosome
endosome lumen
Enrichment
Count
p.adjust
GO-BP
Enrichment

### Figure E9

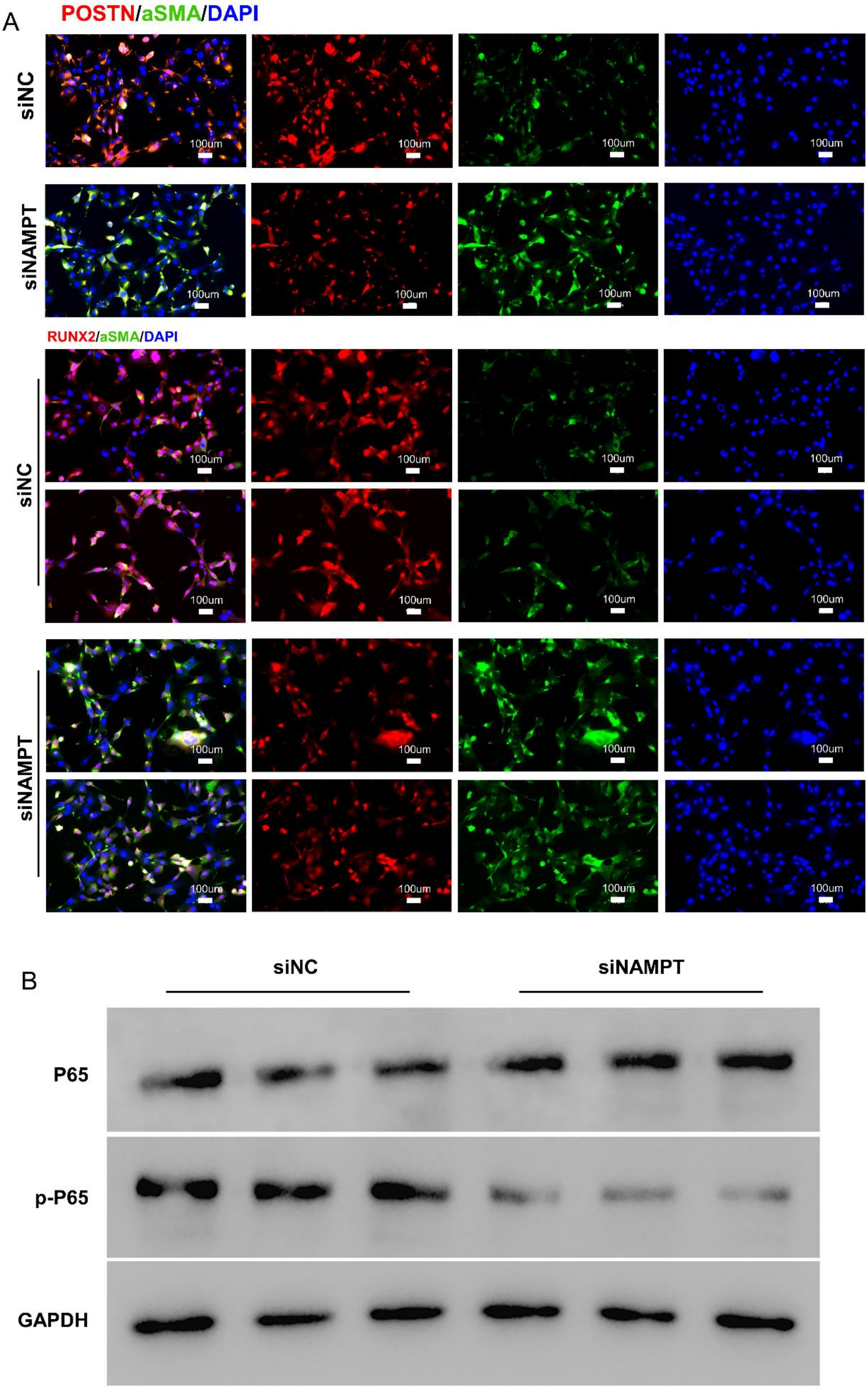
A
POSTN/aSMA/DAPI
siNC
siNAMPT
RUNX2/aSMA/DAPI
siNC
siNAMPT
100um
B
siNC
siNAMPT
P65
p-P65
GAPDH

**Figure E10**
